# Emergence of localized patterns in globally coupled networks of relaxation oscillators with heterogeneous connectivity

**DOI:** 10.1101/100933

**Authors:** Randolph J. Leiser, Horacio G. Rotstein

**Affiliations:** Department of Mathematical Sciences, New Jersey Institute of Technology, Newark, NJ 07102, USA.

## Abstract

Relaxation oscillators may exhibit small amplitude oscillations (SAOs) in addition to the typical large amplitude oscillations (LAOs) as well as abrupt transitions between them (canard phenomenon). Localized cluster patterns in networks of relaxation oscillators consist of one cluster oscillating in the LAO regime or exhibiting mixed-mode oscillations (LAOs interspersed with SAOs), while the other oscillates in the SAO regime. We investigate the mechanisms underlying the generation of localized patterns in globally coupled networks of piecewise-linear (PWL) relaxation oscillators where global feedback acting on the rate of change of the activator (fast variable) involves the inhibitor (slow variable). We also investigate of these patterns are affected by the presence of a diffusive type of coupling whose synchronizing effects compete with the symmetry breaking global feedback effects.

## 1 Introduction

Several chemical, biochemical and biological systems exhibit oscillatory temporal patterns when they are far from equilibrium [1–7]. These phenomena are generated by the nonlinear interplay of positive and negative feedback effects operating at different time scales. Point (single) oscillators require at least one variable (activator) that favors both changes in its own production via autocatalytic effects and the production of a second variable (inhibitor). Inhibitors oppose changes in the activator on a slower time scale. Activators and inhibitors represent different state variables in different systems. Examples are the chemical compounds in the Belousov-Zhabotinsky (BZ) reaction [8,9], the substrate and product in product-activated glycolytic oscillations [4,10], the activator and repressor in genetic oscillators, and the voltage and the recovery variabls of the ionic currents in neurons [5].

In many biologically realistic systems the time scales between activators and inhibitors are well separated, and the resulting oscillations are of relaxation type [2,5]. These are captured by the prototypical van der Pol (VDP) model for a triode circuit [11] and the FitzHugh-Nagumo (FHN) tunnel-diode model for nerve cells [12,13]; and, also, by more detailed models as the Oregonator for the BZ reaction [14–16], the Morris-Lecar model for neuronal oscillations [17] the modified versions of the Selkov model for glycolytic oscillations [18–21] and genetic oscillators [22].

The complexity of individual relaxation oscillators results from the combined effect of two distinct inherent properties: (i) the presence of characteristic types of nonlinearities (typically cubic-like) and (ii) the time scale separation between the participating variables referred to above. In addition to the typical large amplitude oscillations (LAOs) of relaxation type, relaxation oscillators may exhibit small amplitude oscillations (SAOs) with an amplitude difference of roughly an order of magnitude as well as abrupt transitions between them (canard phenomenon) as a control parameter changes through a critical range (exponentially small in the parameter defining the slow time scale) [23–29]. Individual 2D relaxation oscillators may display either SAOs or LAOs, but not both. Higher dimensional relaxation oscillators may exhibit mixed-mode oscillations (MMOs) [30,31], where LAOs are interspersed with SAOs. This creates additional effective time scales.

In addition to the individual oscillators’ intrinsic feedback effects, oscillatory networks have feedback effects that result from the network connectivity and its interaction with the intrinsic properties of the individual oscillators. One such type of feedback is generated by global coupling where each oscillator in the network is affected by the dynamics of the rest through one or more of the participating variables. The effects of global coupling in shaping the network oscillatory patterns has been studied in a variety of systems both experimentally and theoretically. These include oscillatory chemical reactions [32–37], electrochemical oscillators [38–48], laser arrays [49], catalytic reactions [50], salt-water oscillators [51], metabolic oscillators and cellular dynamics [20,52,53], cardiac oscillators [54,55], coupling through quorum sensing [56–60], circadian oscillators [61–63], neuronal networks [5,64–69] and image processing [65,70].

Globally coupled networks of 2D relaxation oscillators have been shown to generate oscillatory cluster patterns [20,32–35,38,39,64,71–75]. Each cluster consists of synchronized in-phase identical oscillators. Oscillators in different clusters differ in at least one of their attributes (e.g., frequency, amplitude and phase). A typical example are the antiphase and, more generally, the phase-locked oscillatory cluster patterns. In a phase-locked two-cluster pattern the two oscillators exhibit LAOs. In some cases, they may also exhibit MMOs, which typically reflect the effects of the network connectivity (e.g., inhibition transiently pushing the activator down or terminating an oscillation before it reaches high enough values), but may also reflect the interaction between the connectivity and the intrinsic canard structure [76] of the individual oscillators [72,73].

A more complex type of pattern that emerges in these globally coupled networks are localized oscillations, where one cluster exhibits LAO or MMOs and the other shows no oscillations or SAOs [32–36,72,73]. Each oscillator in the network is monostable (it can display SAOs or LAOs but not both). The break of symmetry into patterns where each oscillator is in a different amplitude regime requires some type of network heterogeneity such as different cluster sizes or the same cluster size with different global feedback intensities in each cluster. In previous work we showed that the generation of these patterns involves the interplay of the network connectivity and the intrinsic properties of the individual oscillators, particularly their ability to exhibit the canard phenomenon and canard-like SAOs (generated in a supercritical Hopf bifurcation). However, the dynamic mechanisms that give rise to localized oscillatory patterns in networks of relaxation oscillators and how these patterns depend on the properties of the participating oscillators is not fully understood.

The goal of this paper is to address these issues in the context of globally coupled networks where the global feedback acting on the rate of change of the activator involves the inhibitor [32–36,72,73]. An additional goal is to understand how these patterns are affected by the presence of a diffusive type of coupling. Since, in contrast to global inhibition, diffusion tends to synchronize oscillators, their interplay generates a competition between the two opposing effects. We use a cluster reduction of dimensions argument [35] and assume the system is divided into two clusters with the same or different sizes. The effects of the cluster size on the dynamics of this two-cluster networks are absorbed into the global feedback parameter coding for the intensity. Different cluster sizes result in an effective heterogeneous connectivity.

To capture the intrinsic dynamics we use a piecewise-linear (PWL) relaxation oscillator model of FitzHugh-Nagumo (FHN) type, which is an extension of the one we used in [77] to investigate the mechanisms of generation of the canard phenomenon. PWL models can be explicitly analyzed using linear tools of dynamical systems and matching “pieces of solutions” corresponding to consecutive linear regimes. PWL models have been used in a variety of fields as caricature of nonlinear models to provide insights into the dynamics of smooth nonlinear models either to investigate the dynamics of individual nodes or networks [78–104].

As in [77], the activator (*v*) nullcline we use is cubic-like and has four linear pieces (Fig. 1, red curve). The inhibitor (*w*) nullcline is sigmoid-like and has three linear pieces (Fig. 1, green curve). The canard phenomenon requires the presence of the two linear pieces in the middle branch of the *v*-nullcline, but a linear *w*-nullcline is enough. However, localization in models having a linear *w*-nullcline is more difficult to obtain and is less robust than in models having sigmoid-like *w*-nullclines.

**Figure 1:**
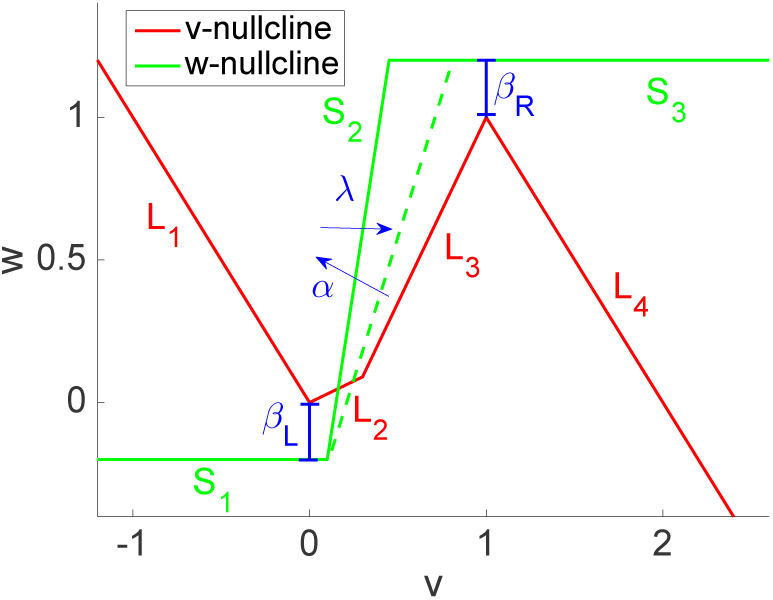
Cubic- and sigmoid-like piecewise linear *v*- and *w*- nullclines for system (1). *The v-nullcline f(v) (red) is given by (2). We used the following parameter values: η = 0.3, v_c_ = 0.3 The w-nullcline g(v; λ) (green) is given by (3). We used the following parameter values for the two superimposed *w*-nullclines: α = 4 (solid-green), α = 2 (dashed-green), λ = 0.3 (solid-green), λ = 0.21 (dashed-green), β_L_ = −0.2, β_R_ = 0.2. The arrows indicate the effects of increasing values of λ and α. Increasing (decreasing) λ displaces the *w*-nullcline to the right (left), while increasing (decreasing) α increases (decreases) the slope of the *w*-nullcline.*

An advantage of using PWL models for this study is that they provide a way of understanding how the intrinsic properties of the individual oscillators affect the network dynamics in terms of the different linear portions of the PWL nullclines whose properties are easily captured by their slopes and end-points. The scenarios we explore in this paper make heavy use of this property. For example, by increasing the values of *β_L_* and *β_R_* in Fig. 1 the *w*-nullcline becomes “more linear” in the region of the phase-plane where the oscillations occur (around the four branches of the cubic-like *v*-nullcine). Along this paper we will compare two such scenarios where the *w*-nullcline is sigmoid-like (as in Fig. 1) and linear-like (relatively large values of both *β_L_* and *β_R_*).

The localized patterns as well as the other types of MMO patterns analyzed in this paper can be a desired or an undesired result of the network activity. For memory devices and working memory [105–108], localized patterns allow for the effective representation of information in the LAO components. In contrast, the presence of localized oscillations may disrupt the communication between neurons [5] and the effective pulsatile secretion of insulin when controlled by glycolytic oscillators or other oscillatory systems (e.g., calcium) [20,109–111] (but see [112]). Our results will contribute to understand the mechanisms underlying the generation of these patterns and how to control or prevent them when necessary.

## 2 Methods

### 2.1 Piecewise linear models of FitzHugh-Nagumo type

We consider the following piecewise linear (PWL) models of FitzHugh-Nagumo (FHN) type

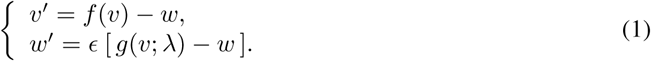

where the prime sign represent the derivative with respect to the variable *t* and the functions *f* and *g* are PWL cubic- and sigmoid-like functions (see Fig. 1) given, respectively, by

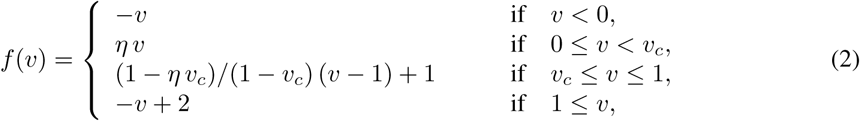

and

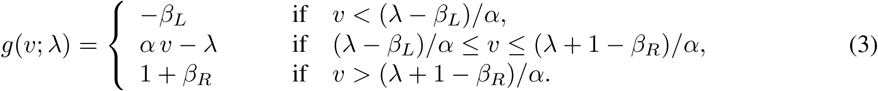

The PWL cubic-like function *f* (Fig. 1, red) has a minimum at (0,0) and a maximum at (1,1). As in the smooth case, this choice ensures that large amplitude oscillations are 𝒪(1) [77]. The parameter *η* governs the slopes of the two middle branches *L*_2_ and *L*_3_. The slope of *L*_3_ also depends on the parameter *v_c_* (*v*-coordinate of the point joining *L*_2_ and *L*_3_). The slopes of both the left (*L*_1_) and right (*L*_4_) branches are equal to −1.

The PWL sigmoid function *g* (Fig. 1, green) has three branches. The two horizontal branches *S*_1_ and *S*_3_ are below and above the minimum and maximum of *f*, respectively. The middle branch *S*_2_ joins these two horizontal branches. The parameter λ controls the displacement of *g* to the right (*lda* > 0) or the left (λ < 0). The parameter *α* controls the slope of the middle branch *S*_2_, which increases with increasing values of *α*. In the limit of *β_L_*, *β_R_* → ∞, the PWL system is the one used in [77] where *g* is a linear function.

### 2.2 Linear regimes and virtual fixed-points

The dynamics of a PWL model of the form (1)-(3) can be divided into four linear regimes *R_k_* (*k* = 1,…, 4), corresponding to the four linear pieces *L_k_* of the cubic-like PWL function *f*(*v*) (Fig. 2). The initial conditions in each regime are equal to the values of the variables *v* and *w* at the end of the previous regime where the trajectory has evolved.

**Figure 2:**
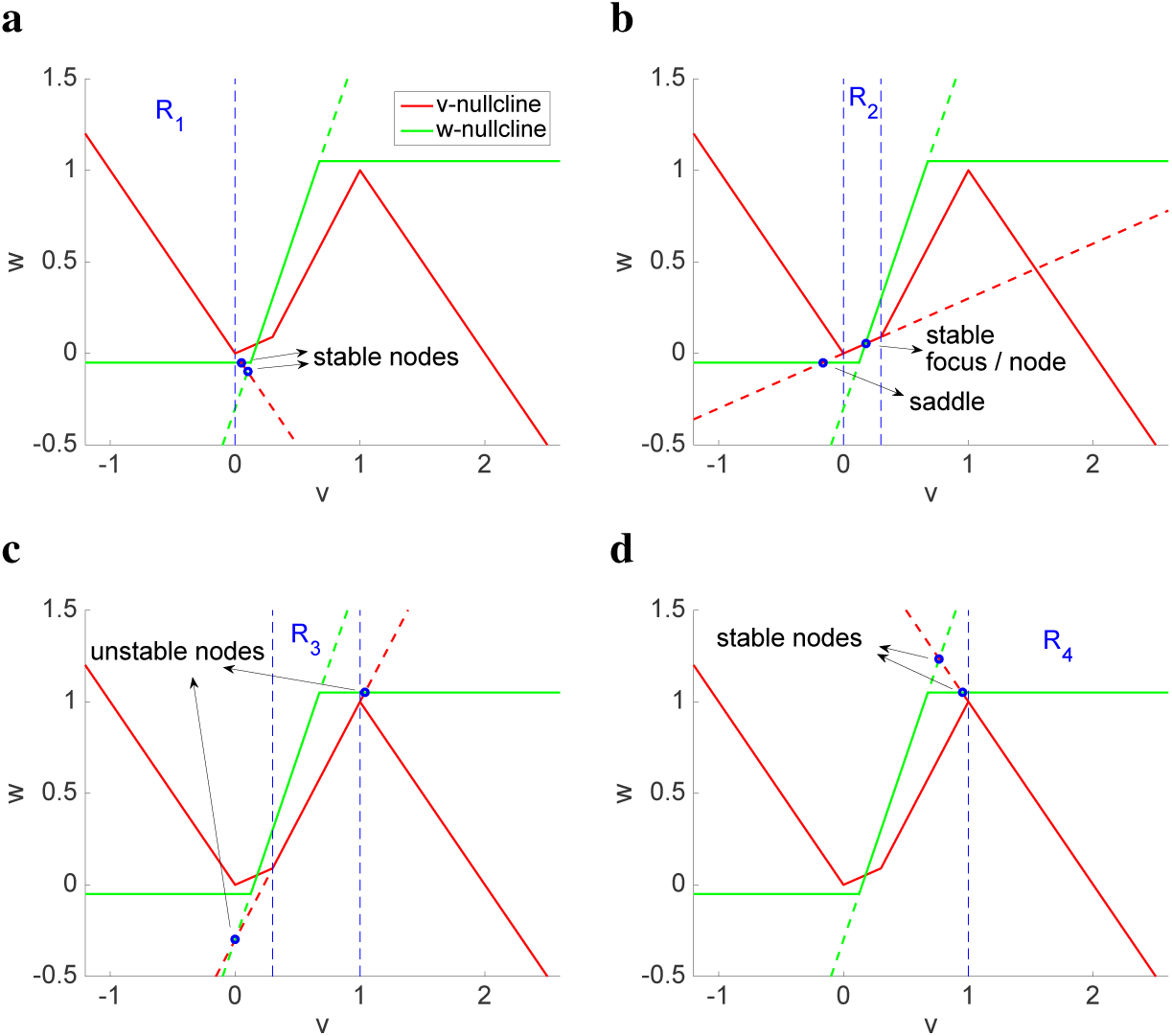
Linear regimes and actual/virtual fixed-point for system (1). *The v-nullcline (red) is as in Fig. 1. For the w-nullcline (green) we used α = 2, λ = 0.3 β_L_ = −0.05 and β_R_ = 0.05. The superimposed dashed-green w-nullcline is linear (extension the linear piece S_2_). The virtual fixed-points for each regime (blue dots) are the intersection between the extensions of the corresponding linear pieces and the w-nullcline. The stable virtual fixed-point for R_2_ coincides with the actual fixed-point.*

In each linear regime the dynamics are organized around a virtual fixed-point (Fig. 2), which results from the intersection between the *w*-nullcline (green line) and the corresponding linear piece (red line) or its extension beyond the boundaries of this regime (dashed-red line). In the latter case the virtual fixed-points do not coincide with the actual fixed-points, and are located outside the corresponding regime, but still play an important role in determining the dynamics in that regime. The trajectories in a given regime never reach the purely virtual stable fixed-points (outside the regime), but their presence provides information about the trajectory’s direction of motion. More specifically, within the boundaries of each regime trajectories evolve according to the linear dynamics defined in that regime as if the dynamics were globally linear, and they “do not feel” that the “rules” governing their evolution will change at a future time when the trajectory moves to a different regime. We refer the reader to [77] for more details.

### 2.3 Networks of PWL oscillators with global inhibitory feedback

We consider networks of PWL oscillators of FHN type of the form (1) globally coupled through the inhibitor variable (*w*)

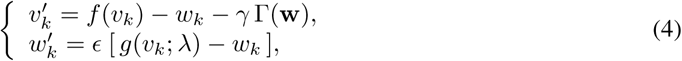

for *k* = 1,…, *N*, where *N* is the total number of oscillators in the network, *γ* ≥ 0 is the global feedback parameter and

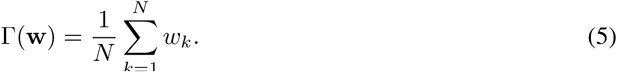

### 2.4 Cluster reduction of dimensions and heterogeneous coupling

Following previous work [35,36,72,73] we assume the network is divided into two clusters where all oscillators in each cluster are identical and have identical dynamics, while oscillators in different clusters may have different dynamics. Accordingly, for a two-cluster network,

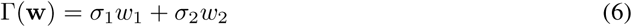

where *σ*_1_ and *σ*_2_ (*σ*_1_ + *σ*_2_ = 1) are the fractions of oscillators in each cluster. Alternatively, the global coupling term (6) can be also interpreted as consisting of clusters with the same fraction of oscillators each, but heterogeneous connectivity.

System (4) with (6) can be written as

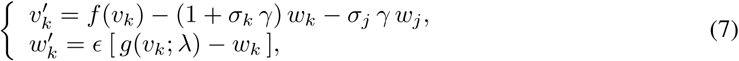

for *k*, *j* = 1, 2 with *j* ≠ *k*.

The zero-level surfaces (“higher-dimensional nullclines”) for the *k^th^* oscillator are given by

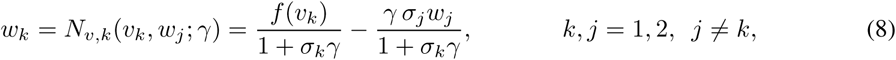

and

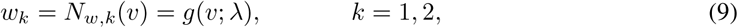

respectively.

Eq. (8) describes a two-dimensional surface having the shape of the first term in the right hand side of *N_v,k_*(*v_k_*, 0; *γ*). For *γ* > 0, we view the nullsurface (8) as the *v*-nullcline for the individual (uncoupled) oscillator *N_v,k_* (*v*, 0; 0), flattened by the effect of the denominator and forced by the second oscillator via the variable *w_j_*(*t*). When there is no ambiguity, we refer to the autonomous part *N_v,k_*(*v_k_*, 0; *γ*) in (8) as the *v*-nullcline for the oscillator *O_k_*. The oscillations in the latter “raise” and “lower” this *v*-nullcline following the dynamics of *w_j_* and therefore affect the evolution of the trajectories in the phase-plane diagrams.

### 2.5 Diffusive coupling between clusters

System (7) with an added diffusion term reads

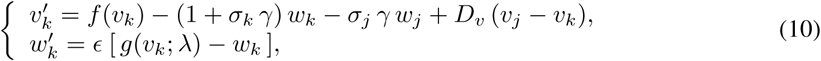

for *k*, *j* = 1,2 with *j* ≠ *k*, where *D_v_* is the diffusion coefficient.

This way of adding diffusion is somehow artificial and does not reflect the diffusive effects in the original system nor is it derived from it. However, its inclusion helps understand the competitive effects of global inhibition and diffusion.

Equation (8) is extended to

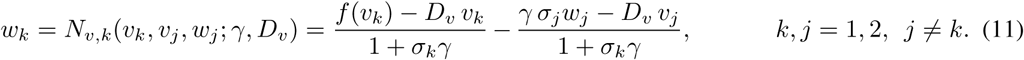

For *D_v_* > 0 the *v*-nullcline *N_v,k_* (*v_k_*, 0,0; *γ*, *D_v_*) is linearly modified by the term *D_v_ v_k_*. In contrast to global coupling, this effect is not homogeneous for all values of *v_k_*, but is dependent on its sign. For positive values of *v_k_* the *v*-nullcline is flattened, while for negative values of *v_k_* the *v*-nullcline is sharpened. The oscillations in *v_j_* “raise” and “lower” this *v*-nullcline following its dynamics. In order for the linear piece *L*_2_ to remain positive for *D_v_* > 0, we will restrict *D_v_* < *η*.

### 2.6 Dynamics of the linear regimes

The dynamics of system (1)-(3) in each linear regime are governed by a system of the form

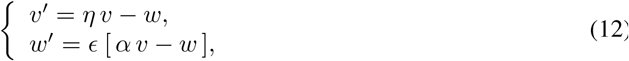

centered at the fixed-point (*v̅*,*w̅*) (virtual or actual) corresponding to each linear regime. In (12) *η* represents the generic slope of each linear piece of *f*(*v*) and not necessarily the slope of the linear piece *L_2_*. The coordinates of the fixed-point for each regime, where *f*(*v*) is described by *η* (*v* − *v̂*) + *ŵ*, are given by

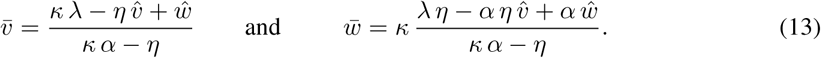

Note that we are using the same notation for the translated system (12) and the original system.

The case *κ* = 1 corresponds to the uncoupled system, while the case *κ* = 1 + *σγ* corresponds to the autonomous part of the globally coupled system (7). The effects of *D_v_* are included in the parameter *η*.

The eigenvalues for each fixed-point are given by

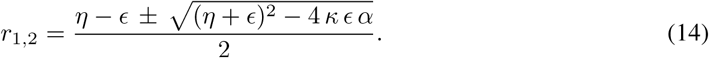

The fixed-points for linear regime (12) are stable if *η* < *∊* and unstable if *η* > *∊*. They are foci if

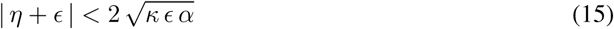

and nodes otherwise. Since *α* ≥ 0 and *κ* > 0, saddles are possible only for *α* = 0. We refer the reader to [77] for a more detailed discussion for the case *κ* = 1. The global feedback parameter *γ* > 0 affects both the location of the fixed-points and the eigenvalues. For large enough values of *γ* a node can transition into a focus.

### 2.7 Numerical simulation

The numerical solutions were computed using the modified Euler method (Runge-Kutta, order 2) [113] with a time step Δ*t* = 0.1 ms (or smaller values of Δ*t* when necessary) in MATLAB (The Mathworks, Natick, MA).

## 3 Results

### 3.1 The canard phenomenon for PWL models of FHN type revisited

In a two-dimensional relaxation oscillator, the canard phenomenon refers to the abrupt transition between small amplitude oscillations (SAOs) and large amplitude oscillations (LAOs) as a control parameter crosses a very small critical range which is exponentially small in the parameter defining the slow time scale [23–29]. We identify this critical range with a critical value for the control parameter (e.g., *λ_c_* if the control parameter is *λ*). If the Hopf bifurcation underlying the creation of the small amplitude limit cycles is supercritical, then the SAOs are stable. In contrast, if the Hopf bifurcation is subcritical, then the small amplitude limit cycle is unstable. The relaxation-type LAOs are always stable. In the subcritical case, there is bistability between a fixed-point and the LAOs for a range of values of the control parameter.

The canard phenomenon for PWL models of FHN type with a linear *w*-nullcline has been described in [77,98] and has been throughly analyzed in [77]. Here we briefly describe it in the context of the PWL models of FHN type with sigmoid-like PWL *w*-nullclines using the parameter λ as the control parameter. Our results are presented in Fig. 3 for two representative types of *w*-nullclines. The first type (*β_L_* = *β_R_* = 0.05 is truly sigmoid-like in the region of the phase-plane where the oscillations occur (Figs. 3-a and -b). The second type (*β_L_* = *β_R_* = 1) is linear in that region (Fig. 3-b). In this paper we refer to them as sigmoid- and linear-like *w*-nullclines, respectively. The primary difference between them is the way they affect the effective time scales in the region of the phase-plane where SAOs and the transition from SAOs to LAOs occur.

**Figure 3:**
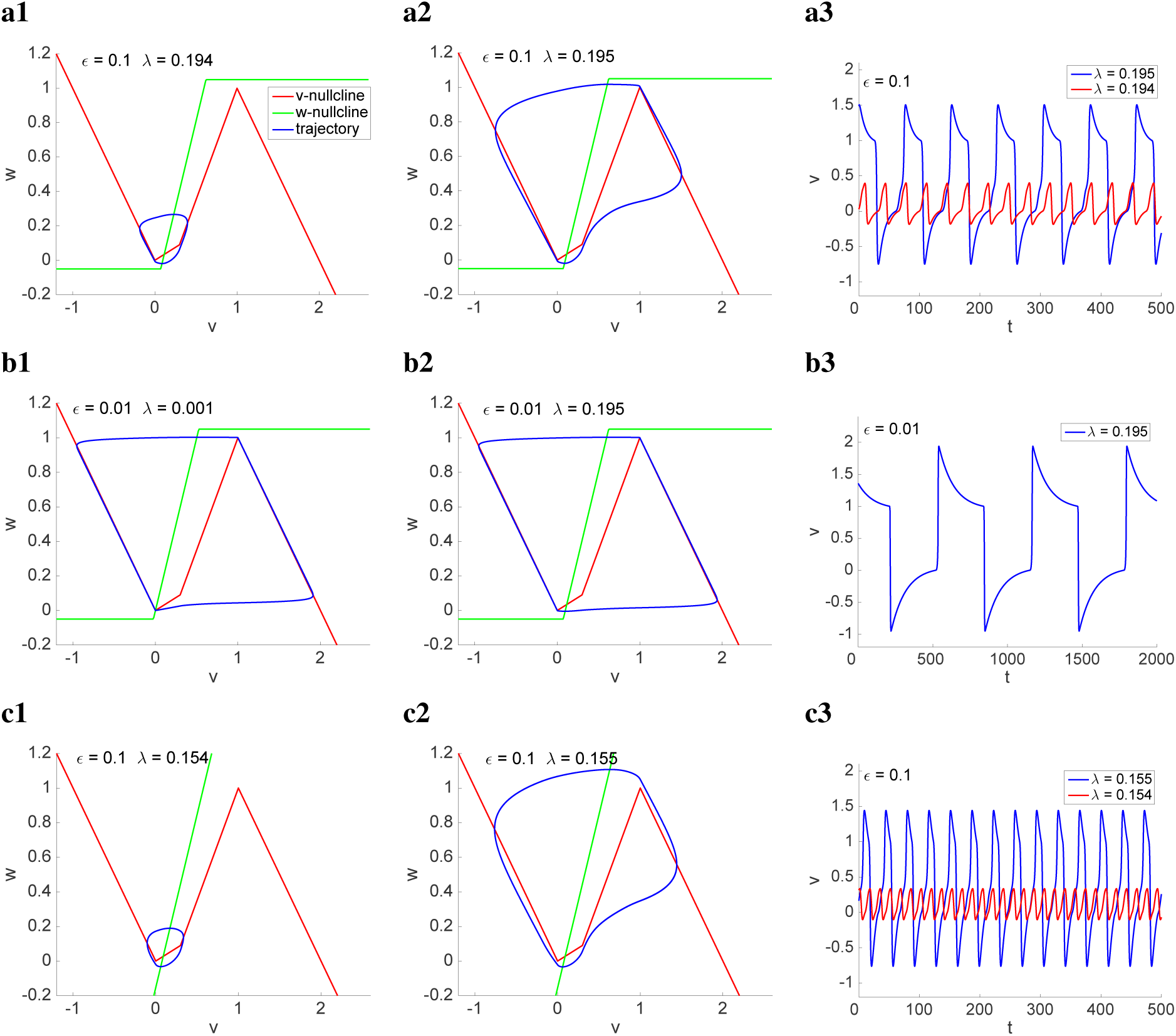
Dynamics of system (1) for representative parameter values. *The v-nullcline (red) is as in Fig. 1. For the w-nullcline we used the following parameter values: α = 2, β_L_ = β_R_ = 0.05 (a, b), β_L_ = β_R_ = 0.5 (c).*

For the SAOs to be generated (Figs. 3-a1 and -c1) the limit cycle must cross either the linear piece *L*_2_ or the first portion of the linear piece *L*_3_ of the *v*-nullcline. Otherwise (Figs. 3-a2 and -c2) the limit cycle trajectory moves into the linear regime *R*_4_ and the system displays LAOs. For a trajectory arriving in *R*_2_ to be able to cross *L*_2_ or the first portion of *L*_3_ the actual fixed-point in *R*_2_ must be a focus (see eq. 15 for *κ* = 1). In addition, the initial amplitude of the trajectory in *R*_2_ (the distance between the actual fixed-point and the initial point in *R*_2_) must be small enough so that the trajectory reaches the *v*-nullcline before reaching the region of fast motion that would cause it to move towards the right branch. For the parameter values in Fig. 3-a, | *η* + *∊* | = 0.4 and 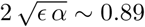 in the linear regime *R*_2_ and therefore the actual fixed-point is a focus. However, as *∊* decreases this inequality may no longer hold. For example, for the parameters in Fig. 3-b, | *η* + *∊* | = 31 and 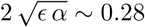) and therefore the actual fixed-point is a node and, as a consequence, the system is no longer able to exhibit the canard phenomenon.

The canard critical value *λ_c_* is affected by the vector field away from the local vicinity of the small amplitude limit cycle (compare Figs. 3-a and -c). The fact that the horizontal branches of the *w*-nullcline in Fig. 3-c are further away from the *v*-nullcline than in Fig. 3-a (*β_L_* and *β_R_* are larger in Fig. 3-c than in Fig. 3-a) causes the oscillation frequency to increase and *λ_c_* to decrease.

### 3.2 The canard phenomenon induced by global feedback

Here we follow previous work [35,36,73] and focus on the dynamics of the one-cluster globally coupled system (7) with *σ*_1_ = 1 (*σ*_2_ = 0). This is not likely to be a realistic situation, but it provides information about how the global feedback affects the oscillatory dynamics and the occurrence of the canard phenomenon in an autonomous system. The results of this section will be helpful in understanding the dynamics of the autonomous part of the two-cluster systems discussed below in this paper.

From (15) with *κ* = 1 + *γ*, increasing values of *γ* (all other parameters fixed) can cause the fixed-point to transition from a node to a focus. In addition, from (14), increasing values of *γ* change the location of the fixed-point. Therefore, the global feedback parameter *γ* can act as the control parameter that induces the canard phenomenon for fixed-values of *λ*.

The top panels in Fig. 4 show curves of the oscillation amplitude versus *γ* for representative parameter values and *α* = 4. The corresponding bottom panels show the phase-plane diagrams for specific values of *γ* (for the blue curves in the top panels). The parameter values in Figs. 4-a and -b are the same, except for *β_L_* and *β_R_* that are smaller in Fig. 4-a than in Fig. 4-b (the *w*-nullcline is sigmoid-like in panel a and linear-like in panel b).

**Figure 4:**
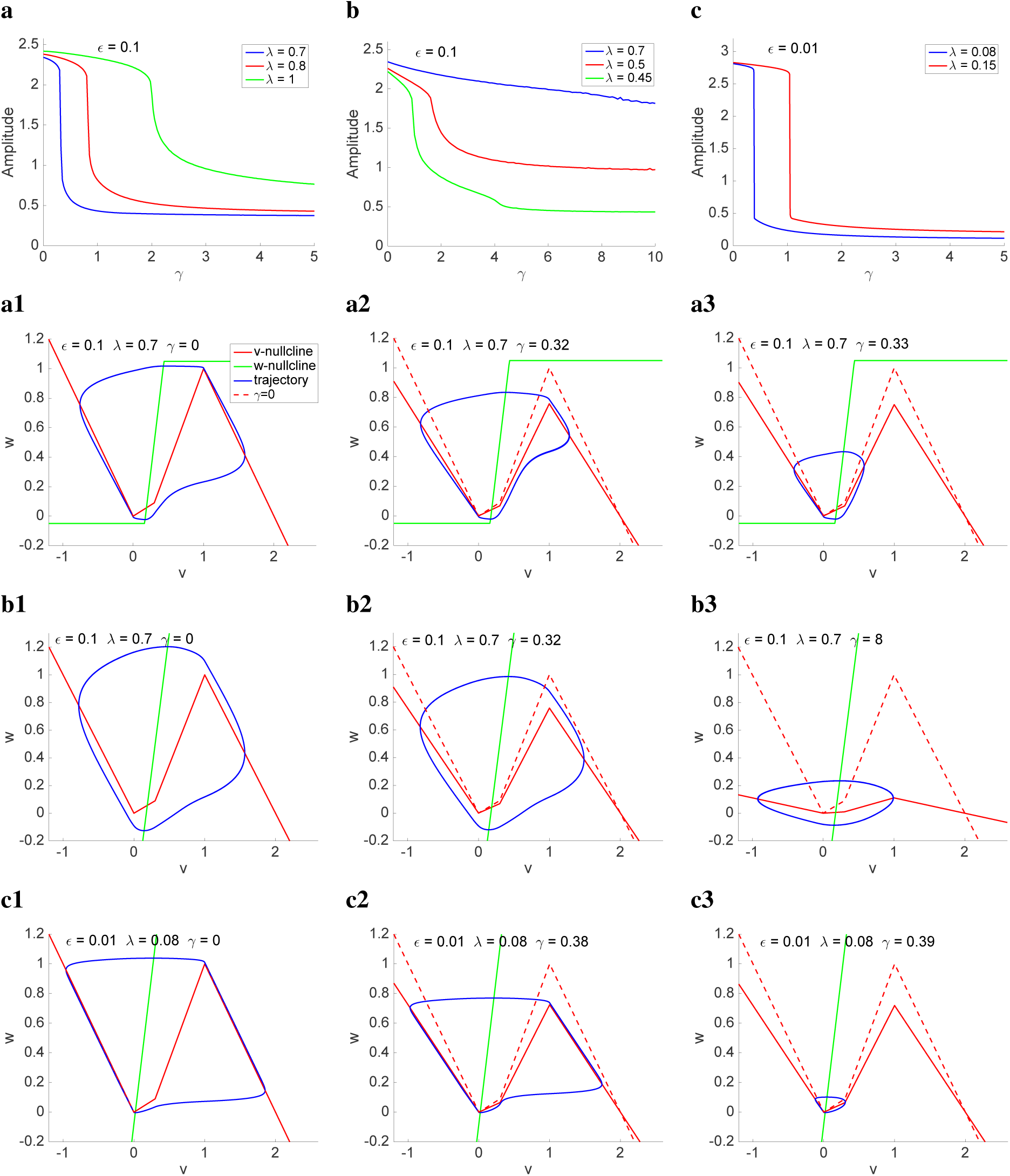
The canard phenomenon induced by global inhibitory feedback in bulk oscillatory systems. *The solid-red v- nullcline corresponds to the actual values of γ used in each panel. The v-nullcline for γ = 0 (dashed-red) is as in Fig. 1 and is presented for reference. For the w-nullcline we used the following parameter values: (**a**) α = 4, ∊ = 0.1, λ = 0.7, β_L_ = β_R_ = 0.05. (**b**) α = 4, ∊ = 0.1, λ = 0.7, β_L_ = β_R_ = 1. (**c**) α = 4, *∈* = 0.01, λ = 0.08, β_L_ = β_R_ = 1.*

For *∊* = 0.1 and the sigmoid-like *w*-nullcline in Fig. 4-a the canard phenomenon is induced by *γ*. The transition is more pronounced for lower values of λ. As *γ* increases, the *v*-nullcline flattens (Figs. 4-a2 and -a3) and for *γ_c_* the limit cycle trajectory is able to cross *L*_3_, thus generating SAOs (Figs. 4-a3), instead of moving towards *R*_4_ to generate LAOs (Figs. 4-a2).

For *∊* = 0.1 and the linear-like *w*-nullcline in Fig. 4-b the system fails to exhibit the canard phenomenon as *γ* increases. The effective time scale separation in the vicinity of the minimum of the *v*-nullcline is smaller than in Fig. 4-a because of the absence of the horizontal piece of the *w*-nullcline (compare panels a1 and a2 with panels b1 and b2), and therefore the limit cycle trajectories are more rounded in Figs. 4-b than in Figs. 4-a. This causes the limit cycle trajectory to move further away from *L*_2_ and *L*_3_ in Fig. 4-b2 than in Fig. 4-a2. As a result, the *v*-nullcline is able to flatten significantly before the limit cycle trajectory is able to cross the middle branch, and therefore the oscillations’ amplitude decreases gradually instead of abruptly. For lower values of λ (red and green curves in Fig. 4-b) the transition from LAOs to SAOs is faster and the final amplitude smaller than for λ = 0.7, but still this transition is not abrupt

A decrease in *∈* for the same parameter values as in Fig. 4-b (including the same linear-like *w*-nullcline) restores the ability of *γ* to induce the canard phenomenon (Fig. 4-c). The decrease in *∊* compensates for the lack of the horizontal pieces of the *w*-nullcline, thus maintaining similar levels of the time scale separation in the vicinity of the minimum of the *v*-nullcline.

As we discussed in the previous section, for *∊* = 0.01 and *α* = 2 the uncoupled oscillator (*γ* = 0) fails to exhibit the canard phenomenon. However, the canard phenomenon can be induced by *γ* (not shown) with similar properties as for *α* = 4. The values of *γ_c_* increase with *λ* and, in contrast to the *α* = 4 case, they are both significantly larger for *α* = 2 than for *α* = 4. Also, the range of values of *γ_c_* spanned by λ is significantly larger for *α* = 2 than for *α* = 4.

### 3.3 Canard and non-canard (standard) SAOs for two interacting oscillators forcing one another

As discussed above, the interaction between two oscillators due to global coupling can be thought of as the oscillators forcing one another through the last term in the first equation in (7). If the product *σ_k_ γ* (*k* = 1,2) is large enough, then the autonomous part of *N_v,k_* (8) can be in an SAO regime. This means that if *w_j_* (*j* = 1, 2 with *j* ≠ *k*) would be artificially made equal to zero, then the oscillator *O_k_* would exhibit the type of canard-like SAOs discussed in the previous section. However, since *w_j_* is not necessarily equal to zero or very small, but also oscillates, then *N_v,k_* raises and shifts down from their baseline location in an oscillatory fashion. This may interfere with the canard SAOs to create the more complex patterns that we discuss in the following sections.

Among these patterns there are the MMOs (LAOs interspersed with SAOs) shown in Fig. 5-a (blue curve). This MMO pattern consists of two types of SAOs. The ones along the ascending phase and the ones on the more shallow phase (Fig. 5-a2). The former correspond to the portion of the trajectory evolving along the left branch of *N*_*v*,1_ (Fig. 5-c) as they respond to the motion of *N*_*v*,1_ following the forcing exerted by *O*_2_ (Fig. 5-c). The second group corresponds to the trajectories moving around the minimum of *N*_*v*,1_ as they are able to cross the linear piece *L*_1_ to create SAOs. We refer to them as canard-like SAOs. The canard-like and standard SAOs are created by different mechanisms. The standard SAOs in *v*_1_ (Fig. 5-a, blue) primarily respond to the oscillatory input from *w*_2_ (Fig. 5-b, red). During the ascending phase, *w*_1_ is decreasing, therefore the oscillations in *v*_2_ and *w*_2_ are intrinsically generated by a canard-like mechanism (Fig. 5-d) that does not require oscillations in the input. The canard-like SAOs are created by the canard-like mechanism described above. Note that although *v*_1_ receives an oscillatory input from *w*_2_, the oscillations in *w*_2_ during the shallow phase have a smaller amplitude than during the ascending phase, indicating that they are less important in the generation of the SAOs in *O*_1_.

**Figure 5:**
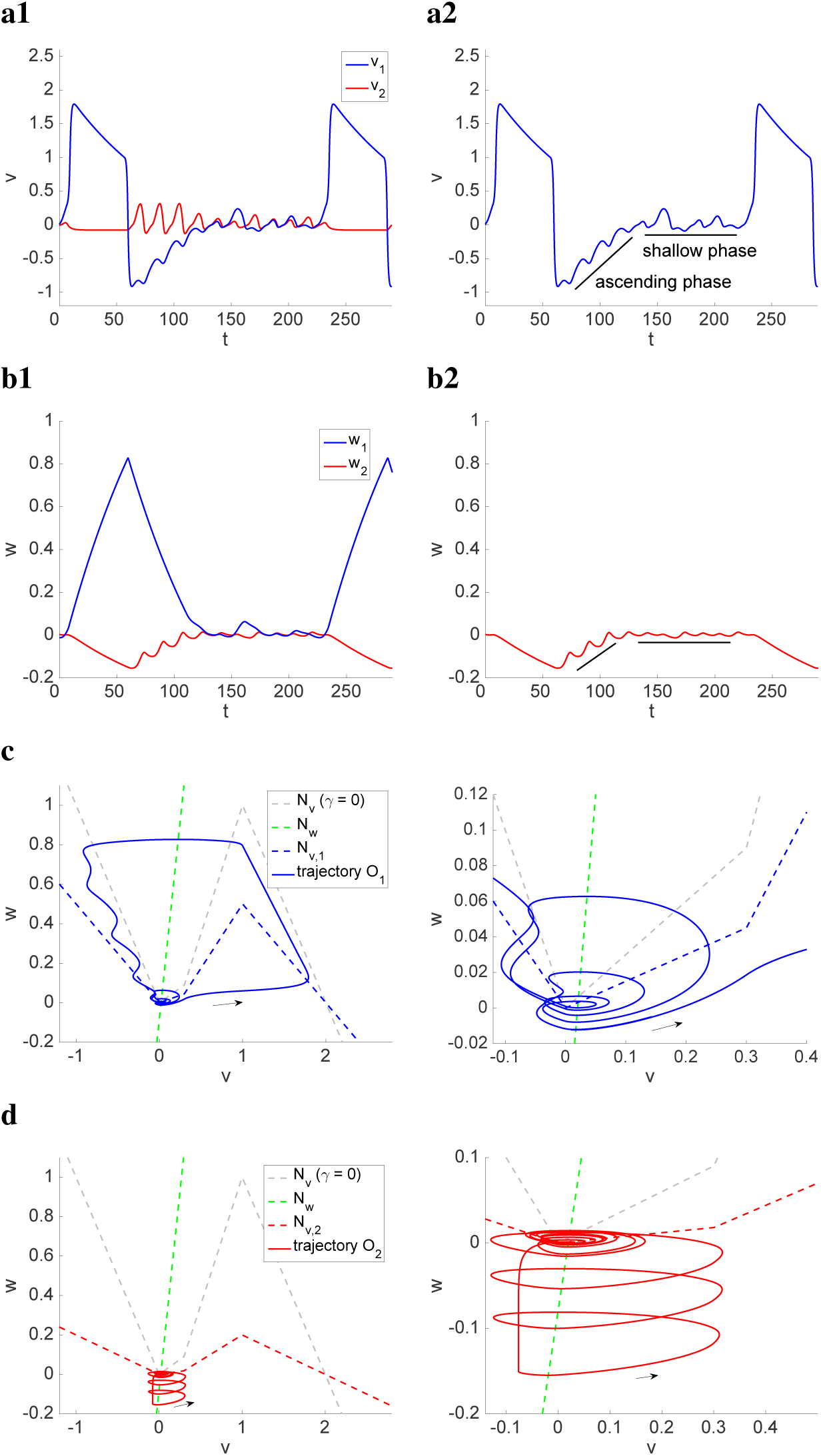
Canard and non-canard (standard) SAOs in two-cluster networks. (**a**) *Curves of v*_1_ *and v*_2_ *vs. t.* (**b**) *Curves of w*_1_ *and w*_2_ *vs. t.* (**c,d**) *Phase-plane diagrams. The gray-dashed curves represent to the vnullcline for the uncoupled system* (*γ* = 0). *The blue- and red-dashed curves represent to the v-nullclines for the autonomous part of the globally coupled system. The green-dashed curves represent the w-nullclines. The solid blue and red curves represent the trajectories of the globally coupled system for the oscillators O*_1_ *and O*_2_ *respectively. The right panels are magnifications of the left ones. We used the following parameter values: α* = 4, *∊* = 0.01, λ = 0.08, *σ*_1_ = 0.2, *σ*_2_ = 0.8, *γ* − 5 *and β_L_* = *β_R_* = 1.

### 3.4 Localized, mixed-mode, phase-locked and SAO network oscillatory patterns

In the next sections we examine the consequences of the global feedback’s ability to induce the canard phenomenon in a single-cluster oscillator (*σ*_1_ = 1 and *σ*_2_ = 0) discussed above for two-cluster network dynamics (*σ*_1_ < 1 and *σ*_2_ > 0). We focus on the case *σ*_1_ =0.2 (*σ*_2_ = 0.8) as a representative case of heterogeneous clusters. As we briefly discuss below, homogeneous clusters (*σ*_1_ = *σ*_2_ = 0.5) produce relatively simple network patterns.

From (7), the autonomous part of each oscillator is affected by both the cluster size (*σ_k_*) and *γ*. In the absence of the forcing exerted by the other oscillator (*w_j_*), the canard phenomenon would be induced by increasing values of both *σ_k_* and *γ* [35,36]. The global feedback parameter critical value for the autonomous part of each oscillatory cluster is given by

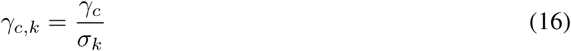

where *γ_c_* is the global feedback parameter critical value for the single-cluster oscillator discussed above (e.g., *γ_c_* = 0.32 in Fig. 4-a and *γ_c_* = 0.38 in Fig. 4-c).

For *σ*_1_ = *σ*_2_ = 0.5, *γ*_*c*,1_ = *γ*_*c*, 2_ and therefore both oscillators would simultaneously be either in the LAO or SAO regime (Fig. 6) with no intermediate types of patterns. However, due to the forcing effects that the oscillators exerts on each other the values of *γ* at which these abrupt transitions occur are larger than the ones predicted by (16). For example, for the parameter values in Fig. 6-a, *γ*_*c*,1_ = *γ*_*c*,2_ ~ 0.76 (see Fig. 4-c) and the transition occurs at *γ* ~ 5.21. For another example, for the parameter values in Fig. 6-b, *γ*_*c*,1_ = *γ*_*c*,2_ ~ 0.72 (not shown) and the transition occurs at *γ* ~ 0.99.

**Figure 6:**
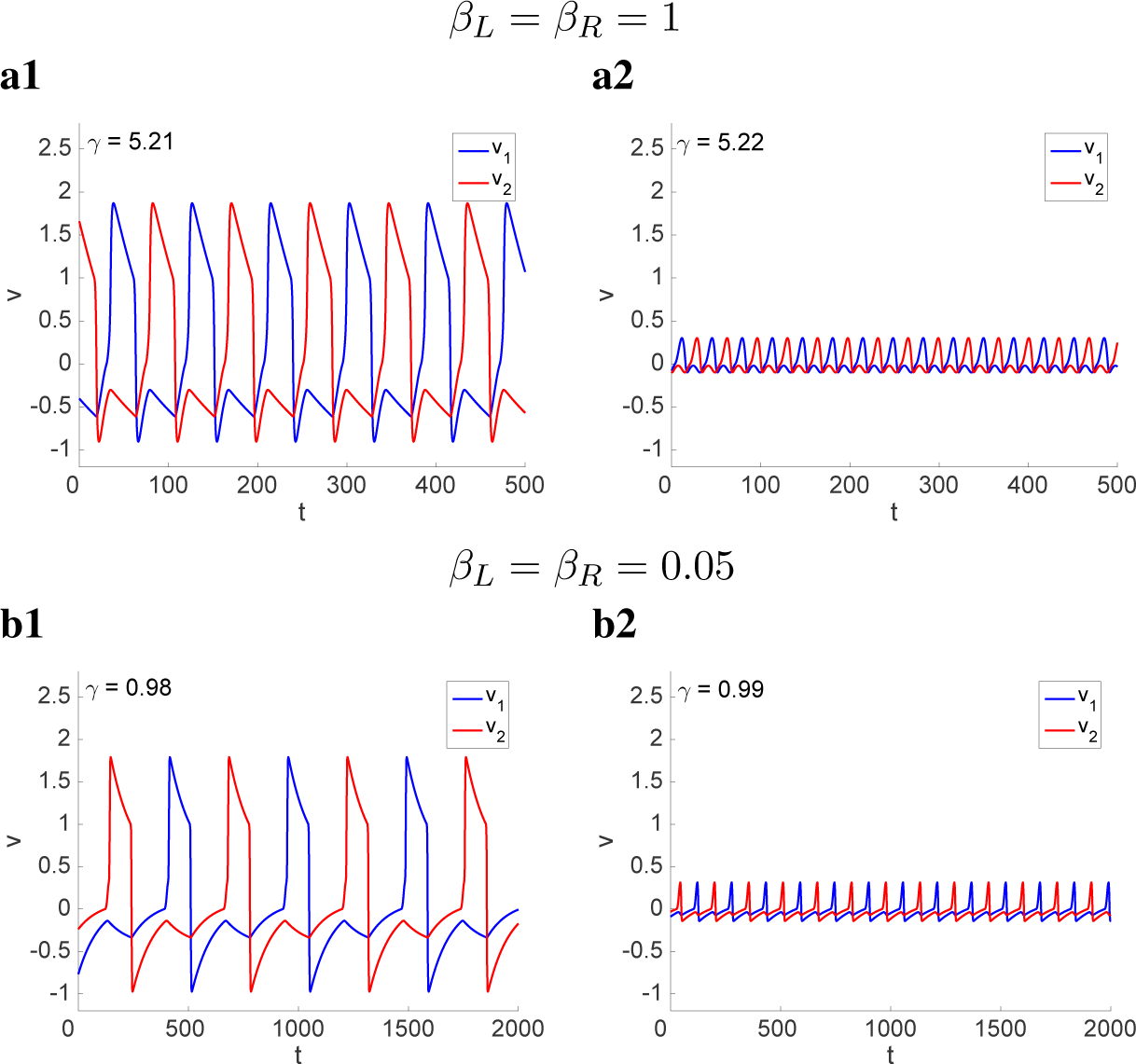
Abrupt transition between antiphase LAO and SAO patterns in two-cluster networks for representative values of γ. (**a**) *Parameter values are as in Fig. 4-c.* (**b**) *Parameter values are as in Fig. 4-c, except for β_L_* = 0.05 *and β_R_* = 0.05 *(same as in Figs. 4-a and -b). We used the following parameter values: α* = 4, *∊* = 0.01, λ = 0.08, *σ*_1_ = *σ*_2_ = 0.5, *β_L_* = *β_R_* = 1 (*a*) and β_L_ = *β_R_* = 0.05 (*b*).

For *σ*_1_ ≠ *σ*_2_ it is be possible for one oscillator (*σ*_1_ < 0.5) to be in the LAO, while the other (*σ*_2_ > 0.5) is in the SAO, thus generating localized patterns (described in more detail below). However, as discussed above, the forcing effects that the oscillators exert on each other is expected to generate more complex dynamics that disrupt this scenario, and it is not a priori clear whether and under what conditions these localized patterns exist. For this to happen, the forcing effects should not interfere with the canard phenomenon for each oscillator. A richer repertoire of intermediate patterns that are not “purely LAO” or “purely SAO” are expected to result from the complex interactions between oscillators as it happens for other systems [72,73].

We have identified various types of network patterns for different parameter regimes.

- Phase-locked LAO patterns (e.g., Figs. 7-a1 and -a2) correspond to both oscillators in the LAO regime. All other parameters fixed, the phase-difference between the two oscillators depends on the relative cluster sizes. For *σ*_1_ = *σ*_2_ = 0.5 the patterns are antiphase (Fig. 6). The underlying mechanisms are qualitatively similar to these described in [72,73] involving the standard SAOs discussed above, and will not be discussed further in the context of this paper.
- Mixed-mode oscillatory (MMO) patterns (e.g., Fig. 7-a3) correspond to either one or both oscillators exhibiting MMOs.
- Localized patterns (e.g., Figs. 7-a5 and -a6 and Figs. 7-b2 and -b3) correspond to one oscillator exhibiting LAOs or MMOs, while the other exhibits exclusively SAOs. From (16), the oscillator with the larger cluster size is the one expected be in the SAO regime.
- LAO localized patterns (e.g., Figs. 10-a3 and -a3) correspond to the two oscillators exhibiting LAOs or MMOs, but the number of LAOs per cycle is different between the two oscillators. The typical situation is one oscillator exhibiting one LAO per cycle, while the other exhibits a burst of LAOs.
- SAO patterns correspond to both oscillators exhibiting SAOs that may or may not be synchronized in phase or have the same amplitude.

**Figure 7:**
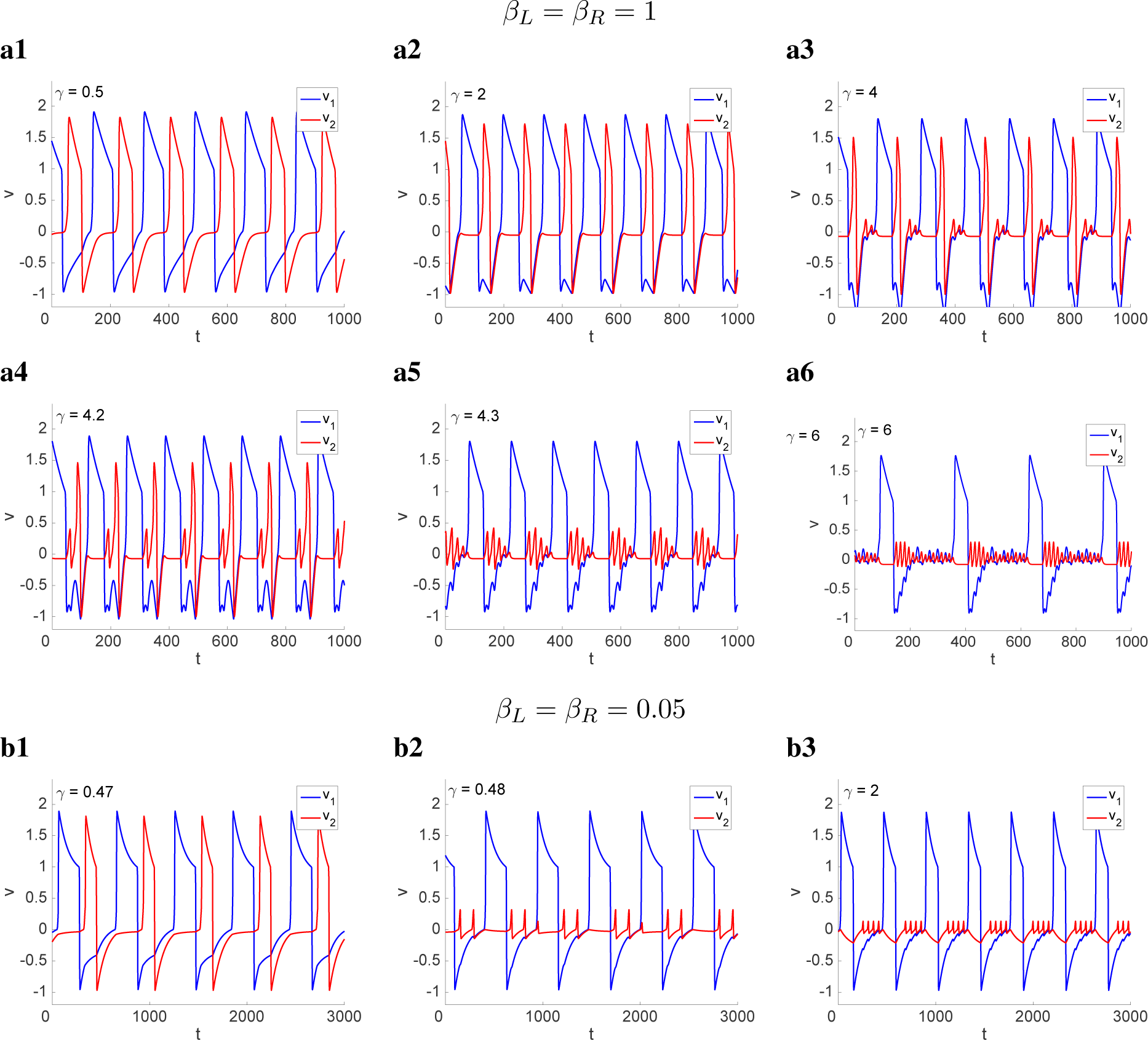
Localization in a two-cluster network for representative values of *γ*. (**a**) *Parameter values are as in Fig. 4-c.* (**b**) *Parameter values are as in Fig. 4-c, except for β_L_* = 0.05 *and β_R_* = 0.05 *(same as in Figs. 4-a and -b). We used the following parameter values: α* = 4, *∊* = 0.01, λ = 0.08, *σ*_1_ = 0.2, *σ*_2_ = 0.8, *β_L_* = *β_R_* = 1 *(a) and β_L_* = *β_R_* = 0.05 *(b)*.

In addition, we have identified various irregular patterns that emerge mostly as transition patterns between these mentioned above. We will not analyze these patterns in this paper.

### 3.5 From phase-locked to localized patterns through network MMOs in the PWL model with a linear-like *w*-nullcline

Fig. 7-a shows various representative two-cluster patterns for the same parameter values as in Fig. 4-c. The global feedback critical values are *γ*_*c*,1_ ~ 1.9 and *γ*_*c*,2_ ~ 0.475. The corresponding phase-plane diagrams are presented in Fig. 8-a.

**Figure 8:**
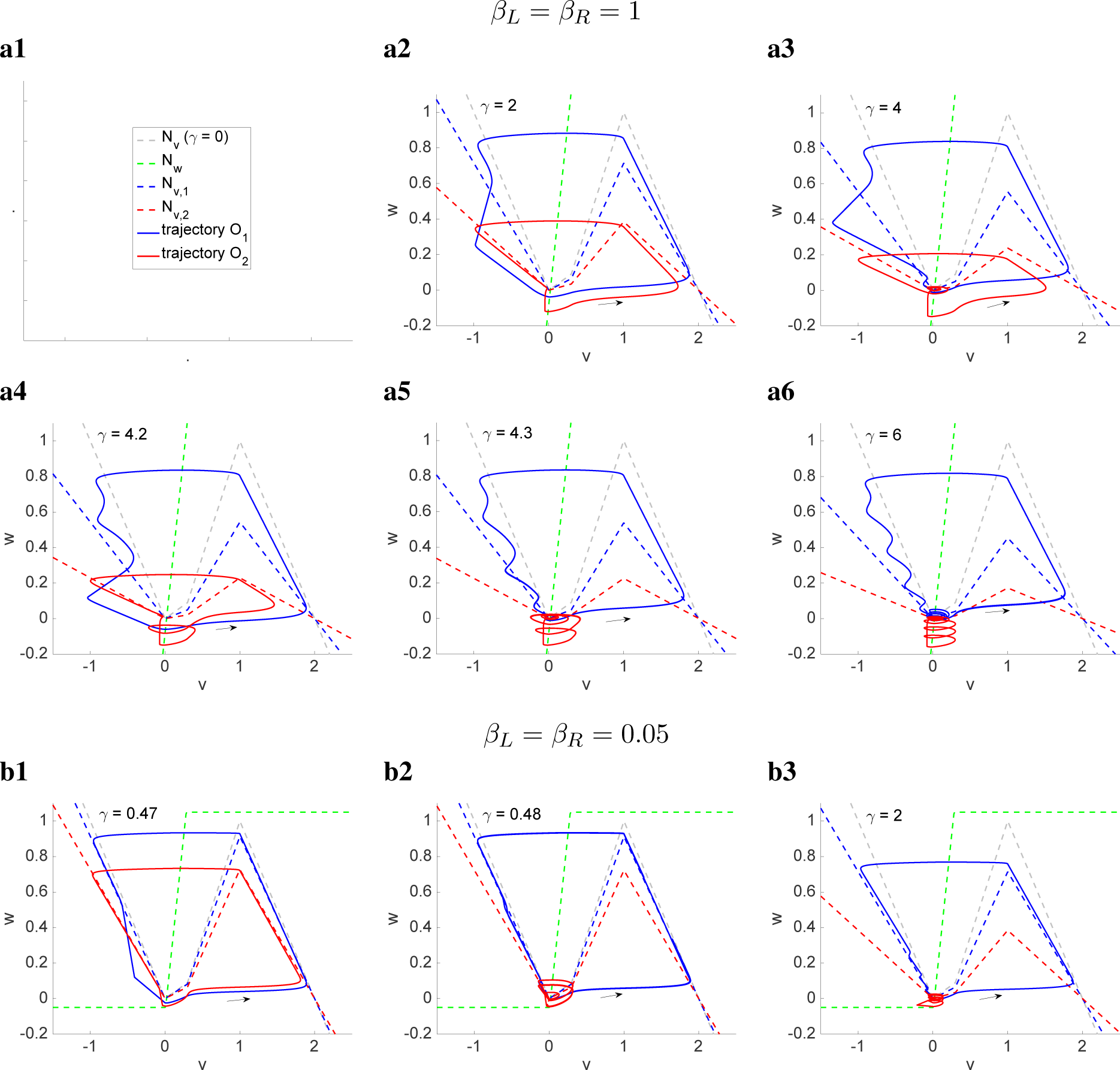
Localization in a two-cluster network for representative values of *γ*. *Phase-plane diagrams for the parameter values is Fig. 7. The gray-dashed curves represent to the vnullcline for the uncoupled system* (*γ* = 0). *The blue- and red-dashed curves represent to the v-nullclines for the autonomous part of the globally coupled system. The green-dashed curves represent the w-nullclines. The solid blue and red curves represent the trajectories of the globally coupled system for the oscillators O*_1_ *and O*_2_ *respectively. We used the following parameter values: α* = 4, *∊* = 0.01, λ = 0.08, *σ*_1_ = 0.2, *σ*_2_ = 0.8, *β_L_* = *β_R_* = 1 *(a) and β_L_* = *β_R_* = 0.05 *(b)*.

For low values of *γ* (Fig. 7-a1 and -a2) the system exhibits phase-locked LAO patterns. The duty cycle is smaller for the larger cluster (oscillator *O*_2_) since its nullcline is flatter (Fig. 8-a2). The relative size of the (smaller to larger) duty cycles for the two oscillators *O*_1_ and *O*_2_ decreases with increasing values of *γ*.

As *γ* increases above these values the system transitions to MMO patterns (Fig. 7-a3 and -a4). The SAOs for *O*_2_ in Fig. 7-a3 are canard-like (Fig. 8-a3) (the limit cycle trajectories cross the linear piece *L*_2_ or at most the early portion of *L*_3_). The last SAO in each cycle for *O*_1_ is also canard-like. They all occur as both *w*_1_ and *w*_2_ are very small so their forcing effects are almost negligible. In contrast, the first SAOs in each cycle are standard (not canard-like) and reflect the motion of *N*_*v*,1_ in response to the dynamics of *O*_2_ as explained in Section 3.3.

During the active phase of *O*_1_, *O*_2_ is almost silent (constant). When *O*_1_ jumps down, *w*_1_ decreases and *N*_*v*,2_ raises, thus releasing *O*_2_. Because *N*_*v*,2_ is flatter than *N*_*v*,1_, *O*_2_ completes the cycle just before *O*_1_, and for some time they are both silent (*w*_1_ ~ 0 and *w*_2_ ~ 0). Fig. 7-a4 corresponds to a slightly higher value of *γ*. This causes the first *O*_2_ oscillation to transition to a SAO. As this happens *O*_1_ is moving along the left branch of *N*_*v*,1_ and continuing to release *O*_2_ from inhibition. As a result, the second *O*_2_ oscillation is a LAO.

For larger values of *γ* the system transitions to localized patterns (Fig. 7-a5 and -a6) where the smaller cluster (*O*_1_) exhibits MMOs and larger cluster (*O*_2_) exhibits canard-like SAOs (Fig. 8-a5 and -a6). The SAOs displayed by *O*_1_ are a combination of canard- and standard SAOs. (as described above) in response to the dynamics of *O*_2_. Note that the transition to localized patterns requires a much larger value of *γ* than the one predicted by *γ*_*c*,1_ and *γ*_*c*,2_.

### 3.6 Abrupt transition between phase-locked LAO and localized patterns for the PWL model with a sigmoid-like *w*-nullcline

Fig. 7-b shows various representative two-cluster patterns for the same parameter values as in Fig. 7-a, but a sigmoid-like *w*-nullcline instead of a linear-like one as in Fig. 7-a. The corresponding phase-plane diagrams are presented in Fig. 8-b. The global feedback critical values are *γ*_*c*,1_ ~ 1.8 and *γ*_*c*,2_ ~ 0.45.

In contrast to the case discussed in the previous section, the transition from phase-locked SAO patterns (Fig. 7-b1) to localized patterns (Fig. 7-b2) is abrupt and occurs for a value of *γ* slightly higher than *γ*_*c*,2_. This is the result of the stronger time scale separation imposed by the sigmoid-like *w*-nullcline, particularly in the regions of the phase-plane where the left and right branches of the *v*-nullcline are located (Fig. 8-b).

When *O*_1_ jumps up, it causes *N*_*v*,2_ to shift down, thus inhibiting *O*_2_. For the parameter values in Fig. 7-b1 (phase-locked SAO patterns), the trajectory for *O*_2_ is above the minimum of *N*_*v*,2_ and it continues to move down along *N*_*v*,2_. After *O*_1_ jumps down and begins to move down along *N*_*v*,1_, decreasing the forcing exerted on *O*_2_, this is released from inhibition and the trajectory moves through *R*_2_ without crossing *L*_2_, thus jumping up.

For the parameter values in Fig. 7-b2 (localized patterns) the trajectory for *O*_2_ is almost at the minimum of *N*_*v*,2_ when *O*_1_ jumps up. The trajectory for *O*_2_ first displays a small non-canard SAO, which is the result of *O*_1_ causing *N*_*v*,2_ to move down, and then two canard SAOs after *O*_1_ jumps down and moves down along *N*_*v*,1_. The larger value of *γ* increases the ability of the trajectory for *O*_2_ to generate canard-like SAOs by crossing *N*_*v*,2_ without jumping up.

The two models considered in this and the previous sections differ in the distances (*β_L_* and *β_R_*) between the horizontal pieces (*S*_1_ and *S*_3_) of the *w*-nullcline and the *v*-nullcline. To determine which one of *β_L_* or *β_R_* has a stronger effect in creating the abrupt transitions between the phase-locked LAO and localized patterns described in this section we looked at models with mixed values of these parameters. We found that for *β_L_* = 1 and *β_R_* = 0.05 the system behaves as in Fig. 7-a, while for *β_L_* = 0.05 and *β_R_* = 1 the system behaves as in Fig. 7-b. This confirms that the increase in the effective time scale separation created by the left horizontal piece of the sigmoid-like w-nullcline is key for the results discussed above (and in the next section).

### 3.7 The oscillatory frequency of the PWL model with sigmoid- and linear-like *w*-nullclines has different monotonic dependencies with *γ*

Comparison between the localized patterns in Figs. 7-a (panels a5 and a6) and -b (panels b2 and b3) shows that the LAO frequency of the oscillator *O*_1_ decreases with increasing values of *γ* for the linear-like *w*-nullcline (Figs. 7-a5 and a6), while it increases with increasing values of *γ* for the sigmoid-like *w*-nullcline (Figs. 7-b2 and b3).

The underlying mechanisms in both cases involve the presence of canard-like SAOs. In Fig. 7-a5, *O*_1_ jumps up right after reaching the minimum of *N*_*v*,1_ (Fig. 8-a5). In Fig. 7-a6, *O*_1_ engages in canard-like SAOs after reaching the minimum of *N*_*v*,1_ (Fig. 8-a6), thus increasing the LAO period. This is the result of the forcing exerted by *O*_2_ and lower time scale separation for the linear-like *w*-nullcline in Fig. 7-a as compared to the sigmoid-like *w*-nullcline in Figs. 7-b.

For the sigmoid-like *w*-nullcline (Figs. 7-b2 and -b3), the number of SAOs per cycle also increases as *γ* increases. However, *O*_1_ jumps up upon reaching the minimum of *N*_*v*,1_. Also, more importantly, the number of cycles per unit of time increases with *γ* because the active phase of *O*_1_ significantly decreases with increasing values of *γ*. This is the result of the flattening of the the *v*-nullcline as *γ* increases and the fact that *O*_1_ jumps down near the maximum of the baseline *N*_*v*,1_.

### 3.8 The localized patterns persist for lower values of *α* for the PWL model with a sigmoid-like *w*-nullcline, but not for a linear-like *w*-nullcline

From our previous discussion about the effects of decreasing values of *α* on the ability of λ and *γ* to induce the canard phenomenon in the uncoupled and coupled systems, respectively, it is not a priori clear whether the localized patterns found in the previous section for *α* = 4 will persist when we decrease *α*. In Fig. 9 we present our results for the same parameter values as in Fig. 7 for *α* = 2 (instead of *α* = 4). For the uncoupled system and *α* = 2 (and all other parameters as in Fig. 9), the PWL model fails to exhibit the canard phenomenon as λ changes. For the one-cluster system, in contrast, changes in *γ* are able induce the canard phenomenon, although for significantly larger values of *γ_c_* than for *α* = 4.

**Figure 9:**
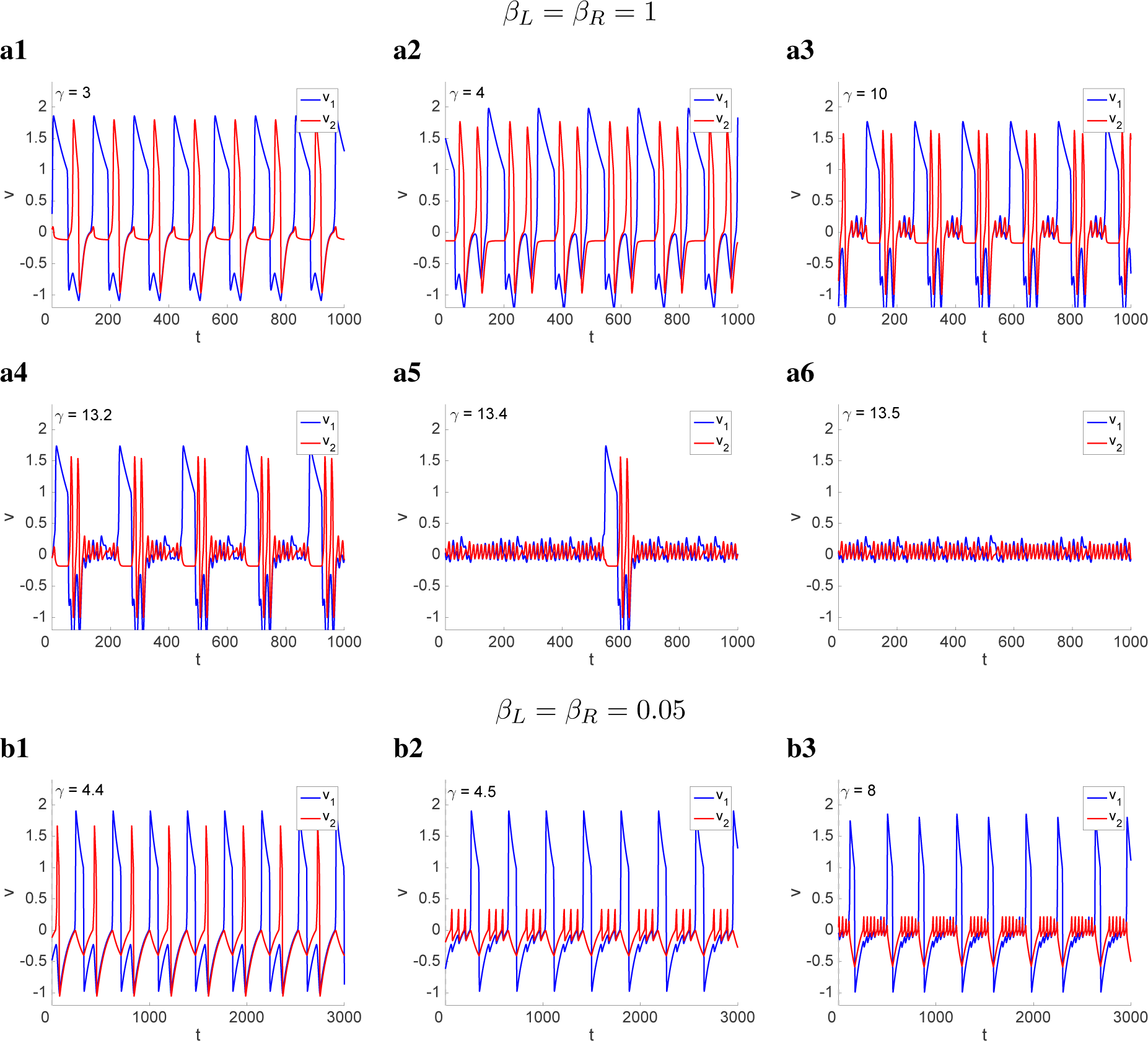
Localized and non-localized patterns in a two-cluster network for representative values of *γ*. *We used the following parameter values: α* = 2, *∊* = 0.01, λ = 0.08, *σ*_1_ = 0.2, *σ*_2_ = 0.8, *β_L_* = *β_R_* = 1 (*a*) *and β_L_* = *β_R_* = 0.05 (*b*).

Our results in Fig. 9-b show that for *α* = 2 and sigmoid-like *w*-nullclines the ability of the PWL model to exhibit abrupt transitions between phase-locked and localized patterns persist with similar properties as for *α* = 4, but the abrupt transition occurs for much higher values of *γ*. In contrast, for linear-like *w*-nullclines the PWL model fails to produce localized patterns (Fig. 9-a). There is an abrupt transition from the LAO patterns in Fig. 9-a4 to the SAO patterns in Fig. 9.

There are additional differences between the patterns in Figs. 9-a and 7-a such as the occurrence of two LAOs per cycle for *α* = 2, which we did not observe for *α* = 4.

### 3.9 The localized patterns are robust to changes in λ for the PWL model with a sigmoid-like *w*-nullcline, but not for a linear-like *w*-nullcline

Increasing values of λ increase the global feedback critical value *γ_c_* (Fig. 4-c), and therefore it increases both *γ*_*c*,1_ and *γ*_*c*,2_, and is expected to increase the values of *γ* at the transition to localized patterns (if they exist) are present. If the values of *γ* are to high, then the *v*-nullcline flattens before the canard phenomenon can be induced by *γ* as for the case illustrated in Fig. 4-b3. Therefore, it is not clear a priori that the transitions observed for λ = 0.08 in Figs. 7 and 8 persist for larger values of *λ*. To address this issue we used the same parameter values as in these figures, but with *λ* = 0.4 (instead of *λ* = 0.08). Our results are presented in Fig. 10.

**Figure 10:**
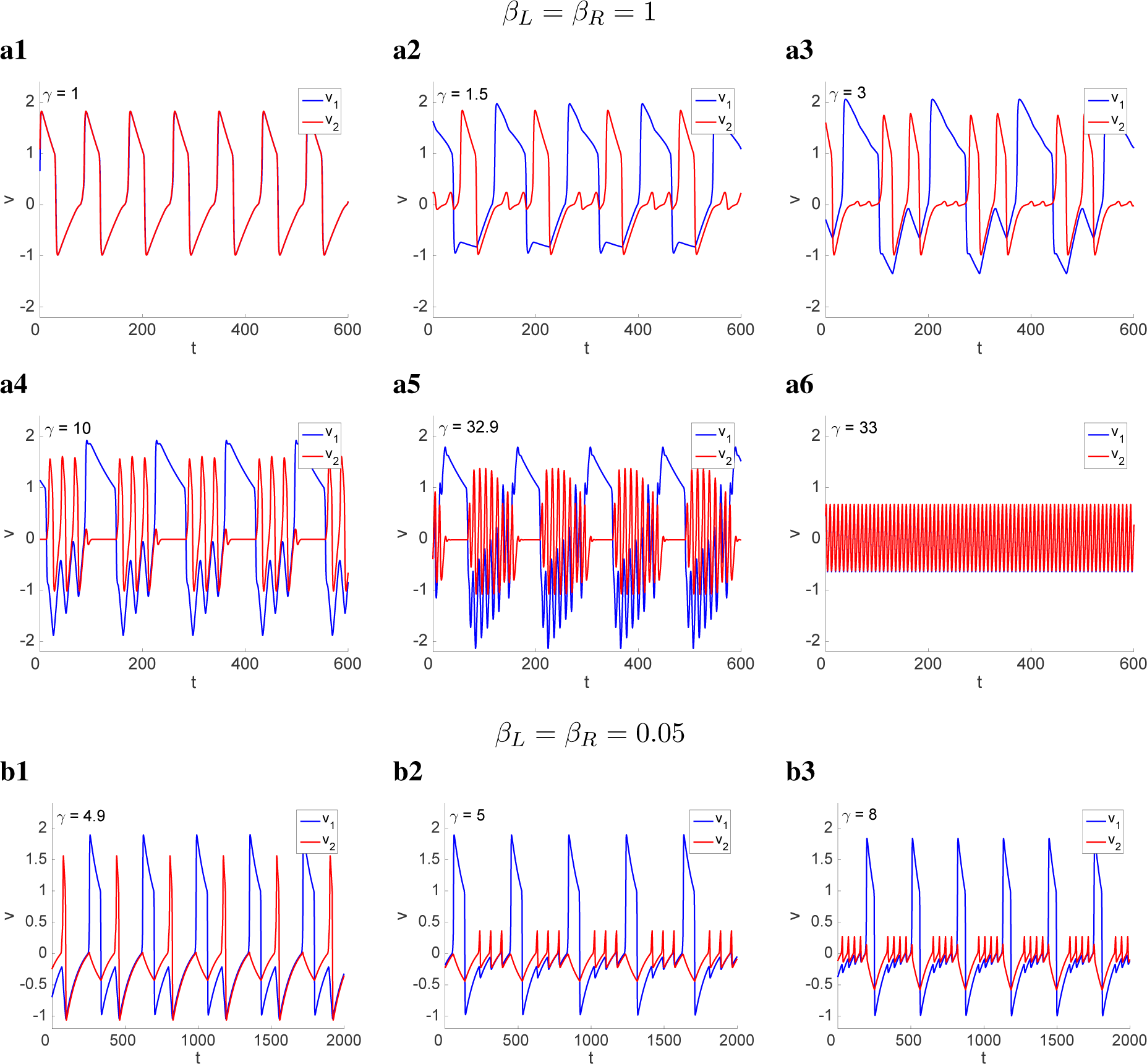
Localized and non-localized patterns in a two-cluster network for representative values of *γ*. *We used the following parameter values: α* = 4, *∊* = 0.01, λ = 0.4, *σ*_1_ = 0.2, *σ*_2_ = 0.8, *β_L_* = *β_R_* = 1 (*a*) *and β_L_* = *β_R_* = 0.05 (*b*).

The model with a sigmoid-like *w*-nullcline (Fig. 10-b) shows an abrupt transition between phase-locked LAOs to localized patterns with similar properties as for λ = 0.08 (Fig. 7-b). In contrast, the patterns displayed for the model with a linear-like *w*-nullcline (Fig. 10-a) differ from these for *λ* = 0.08. Importantly, for *λ* = 0.4 the model does not exhibit localized patterns. Other differences include the presence of in-phase patterns (Fig. 10-a1) and LAO localized patterns (Figs. 10-a3, -a4 and -a5) where the number of LAOs for *O*_2_ per cycle increases with increasing values of *γ*. There is an abrupt transition between these patterns and the ones in Fig. 10-a6.

### 3.10 Localized patterns are more robust for the sigmoid-like *w*-nullcline than for the linear-like *w*-nullcline for higher values of *∊*

In Figs. 11-a and -b we show the different types of patterns for *∊* = 0.1 and the parameter values in Figs. 4-a and -b, respectively. In both cases, for low enough values of *γ* the system shows in-phase patterns (Fig. 11-a1 and -b1), consistently previous findings for the smooth FHN model [72].

**Figure 11:**
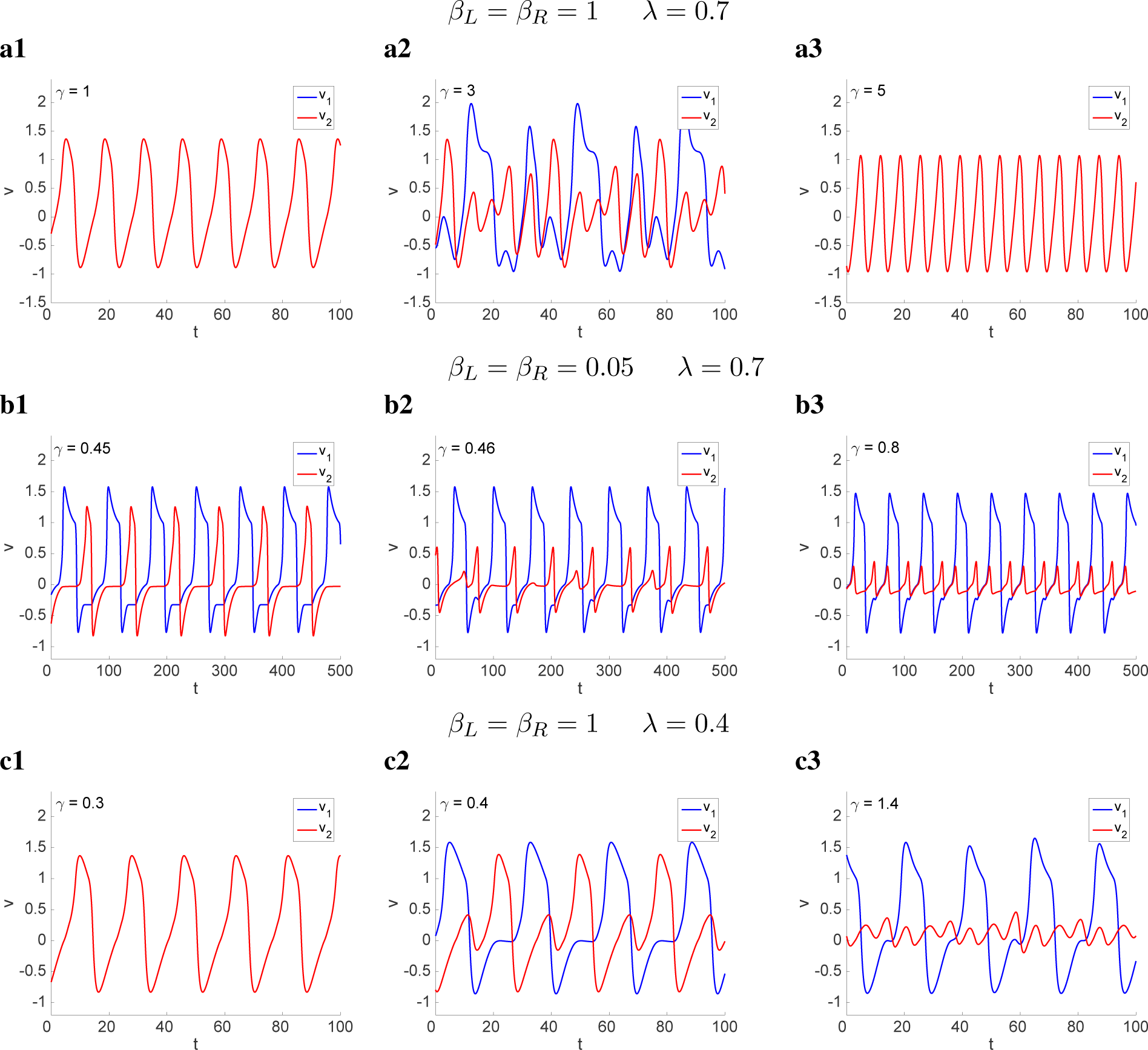
Localization in a two-cluster network for representative values of *γ*. *We used the following parameter values: α* = 4, *∊* = 0.1, λ = 0.7, *σ*_1_ = 0.2, *σ*_2_ = 0.8, *β_L_* = *β_R_* = 1 (*a*) *and β_L_* = *β_R_* = 0.05 (*b*).

As *γ* increases, the patterns in the PWL model with a linear-like *w*-nullcline transition to the complex type of patterns shown in Fig. 11-a2 and then to the synchronized in-phase patterns shown in Fig. 11-a3. The phase-plane diagrams for these patterns (not shown) are qualitatively similar to the ones obtained for the single-cluster case (Fig. 4-b1), which does not exhibit the canard phenomenon as *γ* increases. The absence of localization for the two-cluster system is associated to this lack of ability of the single-cluster system to exhibit the canard phenomenon as *γ* increases.

In contrast, for the PWL model with a sigmoid-like *w*-nullcline, as *γ* increases the patterns transition to the localized patterns shown in Figs. 11-b2 and -b3. The larger and smaller SAOs in Figs. 11-b2 and -b3 correspond to the limit cycle trajectory crossing the linear pieces *L*_3_ and *L*_2_, respectively. In Fig. 11-b3 the limit cycle trajectory crosses *L*_3_ at a lower height than in Fig. 11-b2. A significant difference between these localized patterns and the ones for *∊* = 0.01 (Figs. 7-b and 10-b) is that in the latter the SAOs are interrupted during LAOs, while in the former SAOs and LAOs may occur simultaneously.

While localization does not occur for λ = 0.7 in the PWL model with a linear-like *w*-nullcline, it may be restored for lower values of λ (Fig. 11-c3). For these parameter values the system also shows antiphase patterns (Fig. 11-b2) for lower values of *γ*.

### 3.11 The canard phenomenon can be induced by the diffusion autonomous component

In the next sections we investigate the combined effect of global coupling and diffusion. Here we first look at the effects of the diffusion coefficient *D_v_* on the dynamics of the autonomous part of system (10). This is not a realistic situation, but, as for the effects of *γ* on the one-cluster systems discussed in Section 3.2, it provides information about the dynamics of the autonomous part of each oscillator.

Increasing values of *D_v_* decrease the slopes (*η*) of the linear pieces *L*_2_ and *L*_3_. From (15) this can cause the transition of the actual fixed-point in *R*_2_ from a node to a focus, therefore favoring the occurrence of the canard phenomenon. This is illustrated in Fig. 12 for the same parameter values as in Fig. 4 and *γ* = 0. (The baseline *v*-nullclines for *D_v_* =0 in panels a, b and c, are as in Figs. 4-a1, -b1, and -c1, respectively.)

**Figure 12:**
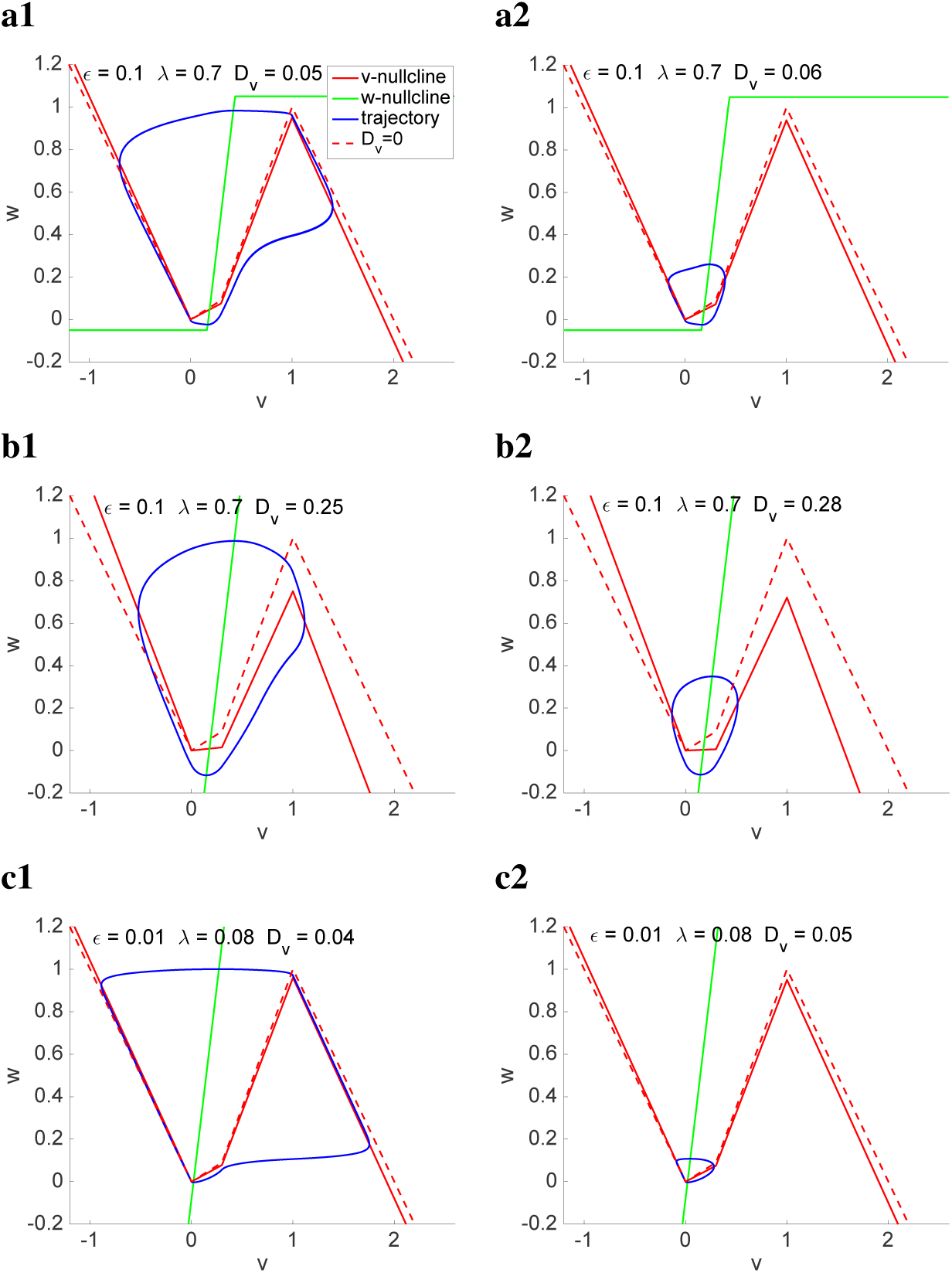
The canard phenomenon induced by diffusion in bulk oscillatory systems. *Parameter values are as in Fig. 4 The baseline v-nullcline for γ* = 0 *(dashed-red) is as in Fig. 1 and is presented for reference. The solid-red v-nullcline corresponds to the actual values of γ used in each panel. For the w-nullcline we used the following parameter values:* (**a**) *α* = 4, *∊* = 0.1, λ = 0.7, *β_L_* = −0.05 *and β_R_* = 0.05. (**b**) *α* = 4, *∊* = 0.1, λ = 0.7, *β_L_* = −1 *and β_R_* = 1. (**c**) *α* = 4, *∊* = 0.01, λ = 0.08, *β_L_* = −1 *and β_R_* = 1.

### 3.12 Interplay of diffusion and global feedback for equal-size clusters: in-phase synchronization and the canard phenomenon

Global feedback and diffusion have opposite effects. While global feedback favors the generation of phase-locked clusters (Fig. 6), diffusion favors in-phase synchronization (Fig. 13-a). Here and in the next section we investigate the patterns that result from the interplay of global coupling and diffusion. For visualization purposes, in Figs. 13 and 14 we present the patterns for the globally coupled system in the absence of diffusion to the left of the dashed-gray line. In a separate set of simulations we have checked that the patterns for both *γ* > 0 and *D_v_* > 0 (right of the dashed-gray line) remain unchanged when global feedback and diffusion are activated simultaneously (not shown).

**Figure 13:**
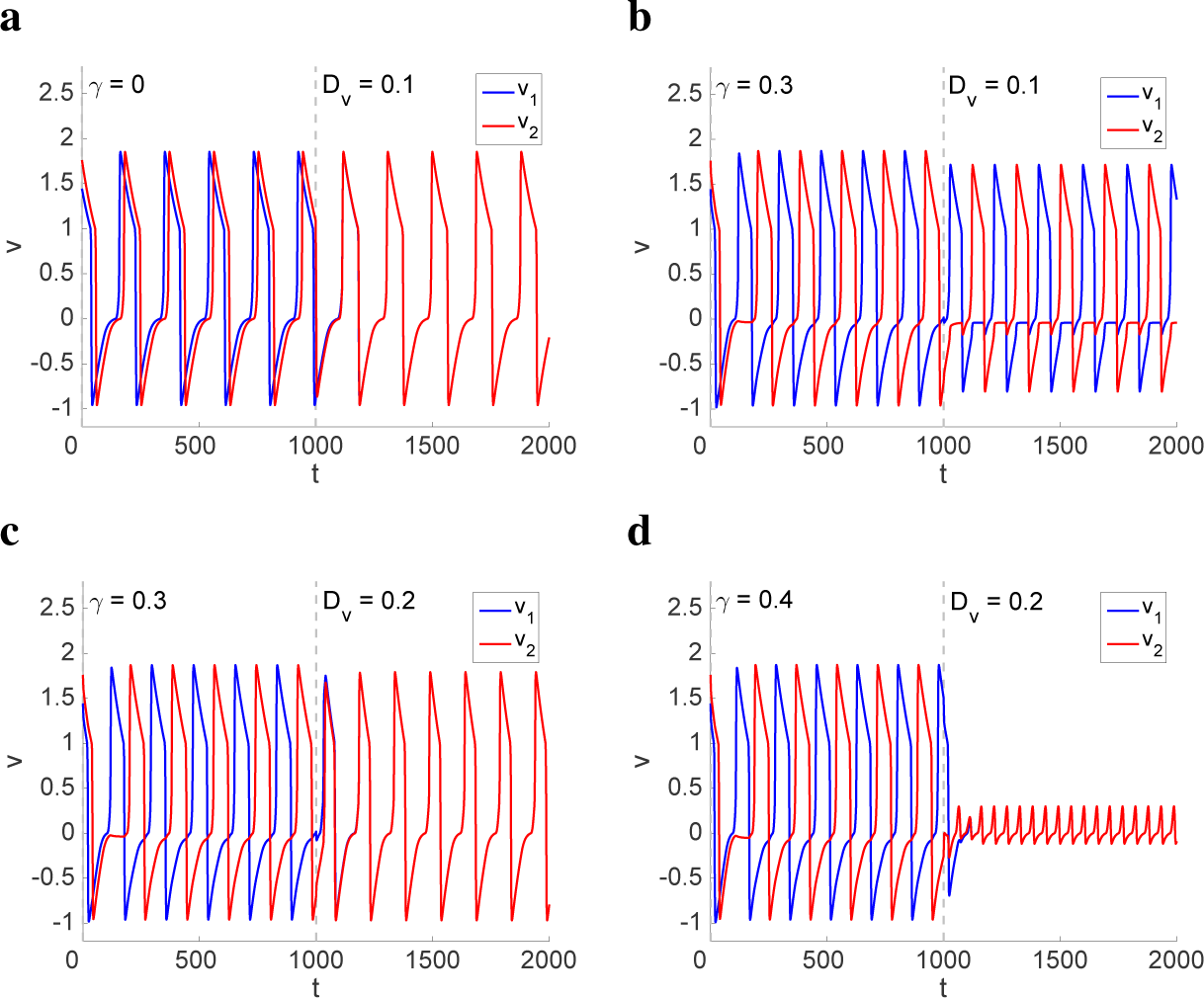
Interplay of global coupling and diffusion in a two-cluster network for representative values of *γ*. *We used the following parameter values: α* = 4, *∊* = 0.01, λ = 0.08, *σ*_1_ = 0.5, *σ*_2_ = 0.5, *β_L_* = *β_R_* = 0.05. *The dashed-gray line indicates the time at which diffusion was activated.*

**Figure 14:**
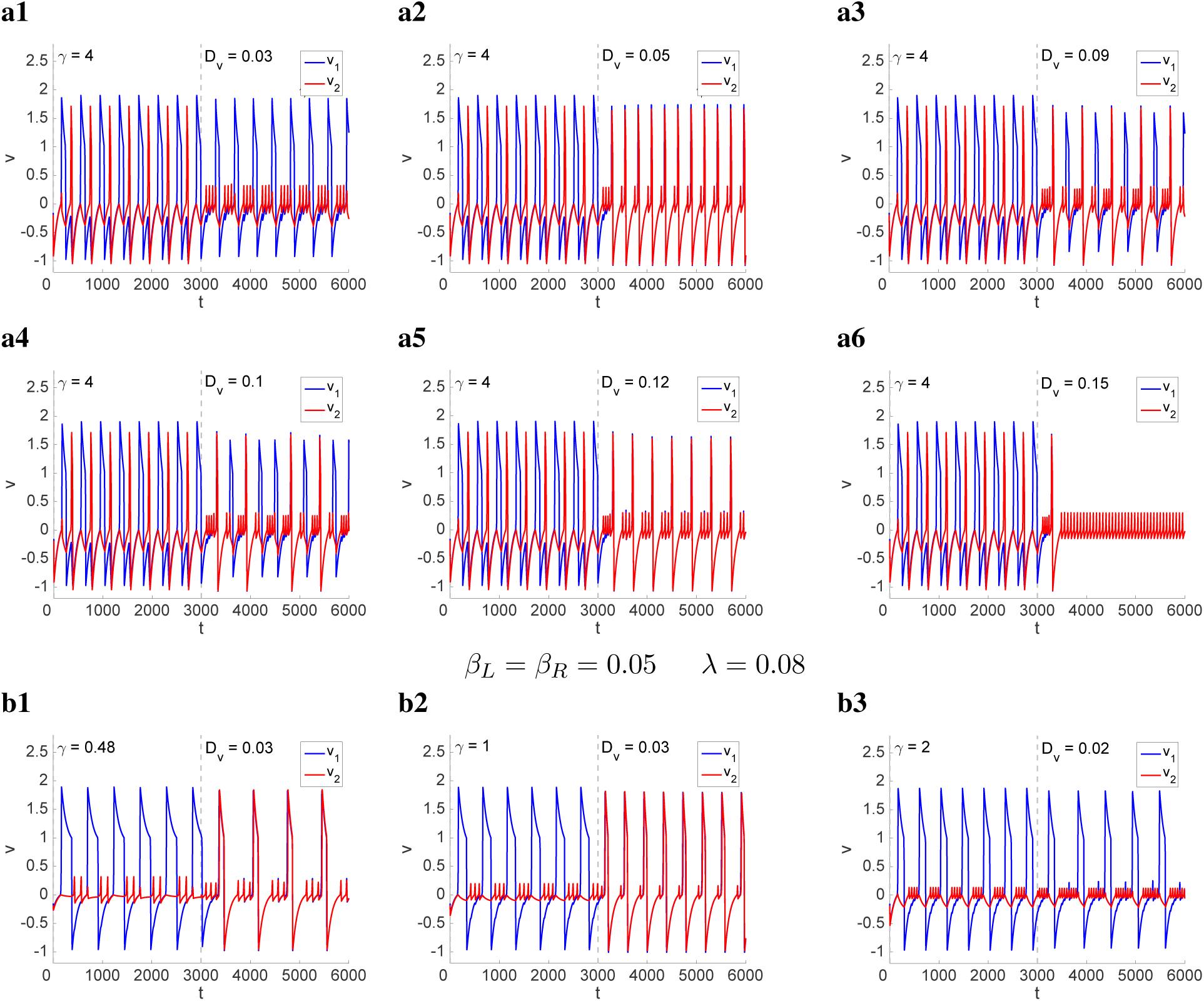
Interplay of global coupling and diffusion in a two-cluster network for representative values of *γ*. *We used the following parameter values: α* = 4, *∊* = 0.01, λ = 0.4 *(a)*, λ = 0.08 *(b)*, *σ*_1_ = 0.2, *σ*_2_ = 0.8, *β_L_* = *β_R_* = 0.05. *The dashed-gray line indicates the time at which diffusion was activated.*

For relative low *D_v_*/*γ* ratios, the system shows antiphase patterns (Fig. 13). As this ratio increases, the two oscillators synchronize in-phase (Fig. 13-c). In-phase patterns are also obtained for *γ* = 0.3 and *D_v_* = 0.15 (not shown) for which *D_v_/γ* = 0.5. For larger values of *γ*, but similar *D_v_*/*γ* ratios, the two oscillators exhibit in-phase SAOs. The increase in *D_v_* does not always cause the transition from LAOs to SAOs since once the two oscillators synchronize in-phase they behave as a single cluster and the diffusive effects are negligible. Therefore, the transition from LAOs to SAOs in these cases depends on whether whether *γ* > *γ_c_* or not. For example, for *γ* = 0.3 and values of *D_v_* larger than the one in Fig. 13-c the patterns remain in the LAO regime in contrast to the patterns in Fig. 13-d).

### 3.13 Interplay of diffusion and global feedback for clusters with different size: Diffusion-induced localized and MMO patterns

In Fig. 14-a1 we illustrate the diffusion-induced localized patterns for the same parameter values as in Fig. 10-b and a relatively low *D_v_*/*γ* ratio. In the absence of diffusion, the system exhibits phase-locked LAOs and localization is induced by increasing values of *γ* (Fig. 10-b). Similar patterns were obtained for other values of *γ* and low *D_v_*/*γ* ratios.

As *D_v_* increases, different types of MMO patterns emerge (Fig. 14-a2 to -a5), which combine the two competing effects of global coupling and diffusion. These patterns include in-phase MMO patterns with different ratios of SAOs and LAOs per cycle (e.g., Fig. 14-a2 and -a5) and MMO patterns where the LAOs in both oscillators are phase-locked (e.g., Fig. 14-a3 and -a4). As *D_v_* increases further, the system exhibits in-phase SAOs.

In Fig. 14-b we show some representative patterns for low *D_v_*/*γ* ratios. Fig. 14-b1 shows a transition from localized to in-phase MMO patterns (similar to this in Fig. 14 -a5) that combine the features of both oscillators when *D_v_* = 0. The in-phase MMOs have a lower LAO frequency than the localized pattern for *D_v_* = 0. Fig. 14-b2 also shows a transition between localized and in-phase MMOs. However, these MMOs have less SAOs per cycle and a higher LAO frequency than for *D_v_* = 0. Finally, in Fig. 14-b3 there is a transition between two types of localized patterns with different ratios of SAOs per cycle and a lower frequency. In all cases, there is relatively abrupt transition between these patterns and SAO patterns, often not synchronized in-phase (not shown). These transitions sometimes involve irregular patterns for very small ranges of *D_v_*

## 4 Discussion

Localized patterns in oscillatory networks where one oscillator (or cluster) exhibits LAOs or MMOs, while the other exhibits SAOs have been observed both experimentally and theoretically [32–36,72,73, 114–116]. In previous work we have established that these type of localized patterns can be obtained in networks of relaxation oscillators such as the FHN model and the Oregonator where the individual oscillators exhibit the supercritical canard phenomenon. In these networks, localized patterns required the presence of heterogeneity in the cluster distribution, which effectively creates heterogeneity in the inter-cluster connectivity. One important aspect of these networks is that the individual oscillators are monostable (they exhibit either LAOs or SAOs, but not both). The break of symmetry in the oscillation amplitude regime between the two (or more) clusters is a network phenomenon. However, how and under what conditions do localized patterns emerge as the result of the interaction between the network connectivity and the intrinsic properties of the individual oscillators (e.g., the canard phenomenon) was not fully understood.

In this paper we set out to address these issues in the context of PWL model of FHN type where the *v*-nullcline is cubic-like and the *w*-nullcline is either sigmoid- or linear-like. This model belongs to the set of minimal models that are able to produce localized patterns. Oscillatory patterns in globally coupled models have also been studied using the so called phase oscillators [37,75,117–122]. In these models, each oscillator is described solely by its phase and the effects of the interaction of oscillators on their amplitude is neglected by assuming weak coupling. These models are successful in capturing the phase-lock cluster patterns where the two oscillators are in the same amplitude regime, but they fail to capture the generation of the more complex patterns that involve more than one oscillatory amplitude regime and transitions between both.

In order to identify the principles that govern how the interplay of the intrinsic properties of the individual oscillators and the network connectivity interact to produce the localized patterns, we have considered a number of representative scenarios which include qualitatively different types of *w*-nullclines (sigmoid- and linear-like) and different parameter values that control the slope of the *w*-nullcline (*α*), its displacement with respect to the *v*-nullcline (λ), and the time scale separation between the participating variables (*∊*).

Our results show that the presence of the supercritical canard phenomenon in the individual oscillatory clusters is a necessary ingredient to produce localized patterns, but it is not sufficient (e.g., Figs. 9-a and 10-a). Localized patterns require a specific tuning between the various model parameters and the shape of the *w*-nullcline. The robustness of these patterns is strongly dependent on the shape of the *w*-nullcline. Models with a sigmoid-like *w*-nullcline produced more robust localized patterns than models with linear-like *w*-nullclines (e.g., Fig. 7) as well as abrupt transitions between phase-locked and localized patterns that were absent in models with linear-like *w*-nullclines.

An additional goal of this study was to understand the effects of the interplay between the two competing types of coupling: global inhibition and diffusion. Global inhibition tends to create clusters. Diffusion is local and tends to cause oscillators to synchronized in-phase. Indeed, when the two clusters have equal size and the oscillators are initially in the LAO regime, the addition of diffusion cause them to synchronize in-phase either in the LAO or SAO regimes depending on the *D_v_*/*γ* ratio. However, when the cluster sizes were different, the addition of diffusion induced localized or MMO network patterns that were either synchronized in-phase or not depending also on the *D_v_*/*γ* ratio. Even when the resulting patterns are synchronized in-phase, they do not resemble the patterns in the absence of diffusion.

We emphasize that the diffusive type of coupling we used in this paper is not realistic and does not reflect the diffusive effects between oscillators in each cluster in the original system. The question of how oscillators in each clusters are held together and how the different cluster sizes are generated as the result of the interplay of global coupling and diffusion remains open.

In this paper we have considered a specific type of global coupling motivated by previous work. Other studies have considered global feedback from the activator variable onto itself, rather than from the inhibitor onto the activator [69,123–126]. More research is needed to establish if and under what conditions localized patterns are possible in these networks and, if they exist, to characterize the similarities and differences between the patterns generated by the two types of global feedback.

An alternative scenario to the one we present here would involve the presence of bistability in the individual oscillators [108]. In this case, the role of the network connectivity would be to separate the oscillators into clusters by causing each oscillator to choose between the stationary solutions of the individual oscillators. This will require the presence of bistability between two oscillatory regimes. Alternatively, the localized solutions would involve one oscillatory and one silent cluster.

Network patterns can be generated by various mechanisms. On one extreme, these patterns can be imposed by the network connectivity with little or no participation of the individual oscillators. Our results highlight the richness of the patterns generated by the interplay of the network connectivity and the intrinsic properties of the individual oscillators.

## 5 Acknowledgments

This work was partially supported by the National Science Foundation grant DMS-1313861 (HGR)

## References

[1] S. H. Strogatz. Nonlinear Dynamics and Chaos. Addison Wesley, Reading MA, 1994.

[2] F. Sagués and I. R. Epstein. Nonlinear chemical dynamics. Dalton Trans., pages 1201–1217, 2003.

[3] J. D. Murray. Mathematical Biology: I. An Introduction. Springer, Berlin, 2002.

[4] J. Keener and J. Sneyd. Mathematical Physiology. Springer-Verlag, New York, 2001.

[5] G. B. Ermentrout and D. Terman. Mathematical Foundations of Neuroscience. Springer, 2010.

[6] A. T. Winfree. The geometry of biological time, 2nd ed. Springer-Verlag, New York, 2001.

[7] I. R. Epstein and J. A. Pojman. An introduction to nonlinear chemical dynamics. Oxford University Press, 1998.

[8] B. P. Belousov. A periodic reaction and its mechanism. Compilation of Abstracts on Radiation Medicine (Med. Publ., Moscow), 147:145, 1959.

[9] A. M. Zhabotinsky. Periodic processes of malonic acid oxidation in a liquid phase. Biofizika, 9:306–311, 1964.

[10] A. Goldbeter. Biochemical oscillations and cellular rhythms: the molecular basis of periodic and chaotic behavior. Cambridge University Press, Cambridge, 1996.

[11] B. van der Pol. On relaxaction oscillations I. Phil. Mag., 2:978–992, 1926.

[12] R. FitzHugh. Thresholds and plateaus in the Hodgkin-Huxley nerve equations. J. Gen. Physiol., 43:867–896, 1960.

[13] J. S. Nagumo, S. Arimoto, and S. Yoshizawa. An active pulse transmission line simulating nerve axon. Proc. IRE, 50:2061–2070, 1962.

[14] R. J. Fields and R. M. Noyes. Oscillations in chemical systems. IV. limit cycle behavior in a model of a real chemical oscillation. J. Chem. Phys., 60:1877–1884, 1974.

[15] B. P. Belousov. A periodic reaction and its mechanism. in Field, R. J. and Burger, M. (Eds.), Oscillations and traveling waves in chemical systems (Wiley, New York), pages 605–613, 1985.

[16] A. M. Zhabotinsky, F. Buchholtz, A. B. Kiyatkin, and I. R. Epstein. Oscillations and waves in metal-ion-catalyzed bromate oscillating reactions in highly oxidized states. J. Phys. Chem., 97:7578–7584, 1993.

[17] H. Morris, C.a nd Lecar. Voltage oscillations in the barnacle giant muscle fiber. Biophysical J., 35:193–213, 1981.

[18] L. Segel and A. Goldbeter. Scaling in biochemical kinetics: dissection of a relaxation oscillator. J. Math. Biol., 32:147–160, 1994.

[19] A. Goldbeter and R. Lefever. Dissipative structures for an allosteric model: application to glycolytic oscillations. Biophys. J., 12:1302–1315, 1972.

[20] D. Gonze, N. Markadieu, and A. Goldbeter. Selection of in-phase or out-of-phase synchronization in a model based on global coupling of cells undergoing metabolic oscillations. Chaos, 18:037127, 2008.

[21] P. Westermark and A. Lansner. A model of the phosphofructokinase and glycolytic oscillations in pancreating β-cell. Biophys. J., 85:126–139, 2003.

[22] R. Guantes and J. Poyatos. Dynamical principles of two-component genetic oscillators. PLoS Computational Biology, 2:e30, 2006.

[23] F. Dumortier and R. Roussarie. Canard cycles and center manifolds. Memoirs of the American Mathematical Society, 121:1–100, 1996.

[24] W. Eckhaus. Relaxation oscillations including a standard chase on French ducks. In Lecture Notes in Mathematics, Springer-Verlag (Berlin Heidelberg), 985:449–497, 1983.

[25] S. M. Baer and T. Erneux. Singular Hopf bifurcation to relaxation oscillations. SIAM J. Appl. Math., 52:1651–1664, 1992.

[26] E. Benoit, J. L. Callot, F Diener, and Diener M. Chasse au Canard, volume 31. Collect. Math., 1981.

[27] M. Krupa and P. Szmolyan. Relaxation oscillation and canard explosion. J. Diff. Eq., 174:312–368, 2001.

[28] M. Krupa and P. Szmolyan. Extending geometric singular perturbation theory to nonhyperbolic points - fold and canard points in two dimensions. SIAM J. Math. Anal., 33(2):286–314, 2001.

[29] F. Dumortier. Techniques in the theory of local bifurcations: Blow-up, normal forms, nilpotent bifurcations, singular perturbations. In Bifurcations and Periodic Orbits of Vector Fields, edited by D. Schlomiuk (Kluwer Academic Press, Dordrecht), pages 19–73, 1993.

[30] M. Brøns, T. J. Kaper, and H. G. Rotstein. Introduction to focus issue: Mixed mode oscillations: Experiment, computation, and analysis. Chaos, 18:015101, 2008.

[31] M. Desroches, J. Guckenheimer, B. Krauskopf, C. Kuehn, H. M. Osinga, and M. Wechselberger. Mixed-mode oscillations with multiple time scales. SIAM Review, 54:211–288, 2012.

[32] V. K. Vanag, L. Yang, M. Dolnik, A. M. Zhabotinsky, and I. R. Epstein. Oscillatory cluster patterns in a homogeneous chemical system with global feedback. Nature, 406:(6794):389–391, 2000.

[33] V. K. Vanag, A. M. Zhabotinsky, and I. R. Epstein. Pattern formation in the Belusov-Zhabotinsky reaction with photochemical global feedback. J. Phys. Chem. A, 104:11566–11577, 2000.

[34] L. Yang, M. Dolnik, A. M. Zhabotinsky, and I. R. Epstein. Oscillatory clusters in a model of the photosensitive Belusov-Zhabotinsky reaction system with global feedback. Phys. Rev. E, 62(5):6414–6420, 2000.

[35] H. G. Rotstein, N. Kopell, A. Zhabotinsky, and I. R. Epstein. A canard mechanism for localization in systems of globally coupled oscillators. SIAM J. Appl. Math., 63:1998–2019, 2003.

[36] H. G. Rotstein, N. Kopell, A. Zhabotinsky, and I. R. Epstein. Canard phenomenon and localization of oscillations in the Belousov-Zhabotinsky reaction with global feedback. J. Chem. Phys, 119:8824–8832, 2003.

[37] A. F. Taylor, P. Kapetanopoulos, B. J. Whitaker, R. Toth, L. Bull, and M. R. Tinsley. Phase clustering in globally coupled photochemical oscillators. Eur. Phys. J. Special Topics, 165:137–149, 2008.

[38] M. Bertram, C. Beta, M. Pollmann, A. S. Mikhailov, H. H. Rotermund, and G. Ertl. Pattern formation on the edge of chaos: Experiments with CO oxidation on a Pt(110) surface under global delayed feedback. Phys. Rev. E, 67:036208, 2003.

[39] M. Bertram and A. S. Mikhailov. Pattern formation on the edge of chaos: Mathematical modeling of CO oxidation on a Pt(110) surface under global delayed feedback. Phys. Rev. E, 67:036207, 2003.

[40] Kim M., M. Bertram, M. Pollmann, A. von Oertzen, A. S. Mikhailov, H. H. Rotermund, and G. Ertl. Controlling chemical turbulence by global delayed feedback: Pattern formation in catalytic CO oxidation on Pt(110). Science, 292:1357–1360, 2001.

[41] F. Plenge, P. Rodin, E. Scholl, and K. Krischer. Breathing current domains in globally coupled electrochemical systems: A comparison with a semiconductor model. Phys. Rev. E, 64:056229, 2001.

[42] F. Plenge, Y.-J. Li, and K. Krischer. Spatial bifurcations in the generic N-NDR electrochemical oscillator with negative global coupling: Theory and surface plasmon experiments. J. Phys. Chem. B, 108:14255–14264, 2004.

[43] N. Baba and K. Krischer. Mixed-mode oscillations and cluster patterns in an electrochemical relaxation oscillator under galvanostatic control. Chaos, 18:015103, 2008.

[44] W. Wang, I. Z. Kiss, and J. L. Hudson. Experiments on arrays of globally coupled chaotic electrochemical oscillators: Synchronization and clustering. Chaos, 10:248–256, 2000.

[45] I. Z. Kiss, W. Wang, and J. L. Hudson. Populations of coupled electrochemical oscillators. Chaos, 12:252–263, 2002.

[46] I. Z. Kiss, Y. Zhai, and J. L. Hudson. Resonance clustering in globally coupled electrochemical oscillators with external forcing. Phys. Rev. Lett., 77:046204, 2008.

[47] V. García-Morales, A. Orlov, and K. Krischer. Subharmonic phase clusters in the complex Ginzburg-Landau equation with nonlinear global coupling. Phys. Rev. E, 82:065202, 2010.

[48] V. García Morales and K. Krischer. Normal-form approach to spatiotemporal pattern formation in globally coupled electrochemical systems. Phys. Rev. E, 78:057201, 2008.

[49] S. Yu. Kourtchatov, V. V. Likhanskii, A. P. Napartovich, F. T. Arecchi, and A. Lapucci. Theory of phase locking of globally coupled laser arrays. Phys. Rev. A, 52:4089–4094, 1995.

[50] M. A. Liauw, P. J. Plath, and N. I. Jaeger. Complex oscillations and global coupling during the catalytic oxidation of CO. J. Chem. Phys, 104:6375–6386, 1996.

[51] K. Miyakawa and K. Yamada. Synchronization and clustering in globally coupled salt-water oscillators. Physica D, 151:217–227, 2001.

[52] Y. X.: Li, J. Halloy, J. L. Martiel, and A. Goldbeter. Suppression of chaos and other dynamical transitions induced by intercellular coupling in a model of cyclic AMP signaling in Dictyostelium cells. Chaos, 2:501–512, 1992.

[53] A. Khadra and Y. X. Li. A model for the pulsatile secretion of gonadotropin-releasing hormone from synchronized hypothalamic neurons. Biophys. J., 91:74–83, 2006.

[54] E. Alvarez-Lacalle, J. F. Rodriguez, and B. Echebarria. Oscillatory regime in excitatory media with global coupling: application to cardiac dynamics. Computers in Cardiology, 35:189–192, 2008.

[55] E. Alvarez-Lacalle and B. Echebarria. Global coupling in excitable media provides a simplified description of mechanoelectrical feedback in cardiac tissue. Phys. Rev. E, 79:031921, 2009.

[56] A. F. Taylor, M. R. Tinsley, F. Wang, Z. Huang, and K. Showalter. Dynanical quorum sensing and synchronization in large populations of chemical oscillators. Science, 323:614–617, 2009.

[57] S. De Monte, F. d’Ovidio, S. Danø, and P. G. Sørensen. Dynamical quorum sensing: Population density encoded in cellular dynamics. Proc. Natl. Acad. Sci. USA, 104:18377–18381, 2007.

[58] J. Wolf and R. Heinrich. Dynamics of two-component biochemical systems in interacting cells; synchronization and desynchronization of oscillations and multiple steady states. BioSystems, 43:1–24, 1997.

[59] J. Wolf, J. Passarge, O. J. Somsen, J. L. Snoep, R. Heinrich, and H. V. Westerhoff. Transduction of intracellular and intercellular dynamics in yeast glycolytic oscillation. Biophys. J., 78:1145–1153, 2000.

[60] J. García-Ojalvo and S. H. Elowitz, M. B. Strogatz. Modeling a synthetic multicellular clock: Repressilators coupled by quorum sensing. Proc. Natl. Acad. Sci. USA, 101:10955–10960, 2004.

[61] C. Liu, D. R. Weaver, S. H. Strogatz, and S. M. Reppert. Cellular construction of a circadian clock: Period determination in the suprachiasmatic nuclei. Cell, 91:855–860, 1997.

[62] D. Gonze, S. Bernard, C. Waltermann, A. Kramer, and H. Herzel. Spontaneous synchronization of coupled circadian oscillators. Biophys. J., 89:107–119, 2005.

[63] L. To, M. A. Henson, E. D. Herzog, and F. J. Doyle III. A molecular model for intercellular synchronization in the mammalian circadian clock. Biophys. J., 92:3792–3803, 2007.

[64] D. Golomb and J. Rinzel. Clustering in globally coupled inhibitory neurons. Physica D, 72:259282, 1994.

[65] D. Terman and Wang D. L. Global competition and local cooperation in a network of neural oscillators. Physica D, 81:148–176, 1995.

[66] J. Rubin and D. Terman. Analysis of clustered firing patterns in synaptically coupled networks of oscillators. J. Math. Biol., 41:513–545, 2000.

[67] D. M. Durand, E.-H. Park, and A. L. Jensen. Potassium diffusive coupling in neural networks. Philos. Trans. R. Soc. B, 365:2347–2362, 2010.

[68] M. G. Rosenblum and A. S. Pikovsky. Controlling synchronization in an ensemble of globally coupled oscillators. Phys. Rev. Lett., 92:114102, 2004.

[69] R. A. Stefanescu and V. K. Jirsa. A low dimensional description of globally coupled heterogeneous neural networks of excitatory and inhibitory neurons. PLoS Computational Biology, 4:e1000219, 2008.

[70] D. Wang. Relaxation oscillators and networks. In Encyclopedia of Electrical and Electronics Engineering, J. Webster (ed.), pages 1–10, 2007.

[71] M. Bertram and A. S. Mikhailov. Pattern formation in a surface chemical reaction with global delayed feedback. Phys. Rev. E, 63:066102, 2001.

[72] H. G. Rotstein and H. Wu. Swing, release, and escape mechanisms contribute to the generation of phase-locked cluster paterns in a globally coupled Fitzhugh-Nagumo model. Phys. Rev. E, 86:066207, 2012.

[73] H. G. Rotstein and H. Wu. Dynamic mechanisms of generation of oscillatory cluster patterns in a globally coupled chemical system. J. Chem. Phys., 137:104908, 2012.

[74] D. Golomb, X.-J. Wang, and J. Rinzel. Synchronization properties of spindle oscillations in a thalamic reticular nucleus model. J. Neurophysiol., 72:1109–1126, 1994.

[75] D. Golomb, D. Hansel, B. Shraiman, and H. Sompolinksy. Clustering in globally coupled phase oscillators. Phys. Rev. A, 45:3516–3530, 1992.

[76] H. G. Rotstein, T. Oppermann, J. A. White, and N. Kopell. A reduced model for medial entorhinal cortex stellate cells: Subthreshold oscillations, spiking and synchronization. J. Comp. Neurosci., 21:271–292, 2006.

[77] H. G. Rotstein, S. Coombes, and A. M. Gheorghe. Canard-like explosion of limit cycles in two-dimensional piecewise-linear models of FitzHugh-Nagumo type. SIAM J. Appl. Dyn. Systems, 11:135–180, 2012.

[78] H. P. McKean. Nagumo’s equation. Advances in Mathematics, 4:209–223, 1970.

[79] W. P. Wang. Multiple impulse solutions to McKean’s caricature of the nerve equation. I. Existence. Communications on pure and applied mathematics, 41:71–103, 1988.

[80] W. P. Wang. Multiple impulse solutions to McKean’s caricature of the nerve equation. II. Stability. Communications on pure and applied mathematics, 41:997–1025, 1988.

[81] A. Tonnelier. The McKean’s caricature of the FitzHugh-Nagumo model I. The space-clamped system. SIAM Journal on Applied Mathematics, 63:459–484, 2002.

[82] J. Rinzel and J. B. Keller. Traveling wave solutions of a nerve conduction equation. Biophysical J., 13:1313–1337, 1973.

[83] J. Rinzel. Neutrally stable traveling wave solutions of nerve conduction equations. J. Math. Biol., 2:205–217, 1975.

[84] J. Rinzel. Spatial stability of traveling wave solutions of a nerve conduction equation. Biophysical J., 15:975–988, 1975.

[85] S. Coombes and A. H. Osbaldestin. Period adding bifurcations and chaos in a periodically stimulated excitable neural relaxation oscillator. Physical Review E, 62:4057–4066, 2000.

[86] A. Tonnelier and W. Gerstner. Piecewise linear differential equations and integrate-and-fire neurons: Insights from two-dimensional membrane models. Physical Review E, 67: 021908(1-16), 2003.

[87] S. Coombes. Neuronal networks with gap junctions: A study of piece-wise linear planar neuron models. SIAM Journal on Applied Dynamical Systems, 7:1101–1129, 2008.

[88] S. J. Hogan. On the dynamics of rigid-block motion under harmonic forcing. Proceedings of the Royal Society of London. Series A, 425:441–476, 1989.

[89] S. H. Doole and S. J. Hogan. Non-linear dynamics of the extended Lazer-McKenna bridge oscillation model. Dynamics and Stability of Systems, 15:43–58, 2000.

[90] M. di Bernardo, C. Budd, A. R. Champneys, and P. Kowalczyk. Piecewise-smooth Dynamical Systems: Theory and Applications. Springer, 2007.

[91] G. M. Maggio, M. di Bernardo, and M. P. Kennedy. Nonsmooth bifurcations in a piecewise-linear model of the Colpitts oscillator. IEEE Transactions on Circuts and Systems-I: Fundamental Theory and Applications, 47:1160–1177, 2000.

[92] A. F. Filippov. Differential Equations with Discontinuous Righthand Sides. Kluwer Academic Publishers, Dordrecht, 1988.

[93] E. Plahte and S. Kjoglum. Analysis and generic properties of gene regulatory networks with graded response functions. Physica D, 201:150–176, 2005.

[94] F. Grognard, H. de Jong, and J.-L. Gouze. Biology and Control Theory: Current Challenges. Lecture Notes in Control and Information Science (LNCIS) 357 (I Queinnec and S Tarbouriech and G Garcia and S Niculescu, Eds.). Springer, 2007.

[95] J. Llibre, E. Nu nez, and A. E. Teruel. Limit cycles for planar piecewise linear differential systems via first integrals. Qualitative theory of dynamical systems, 3:29–50, 2002.

[96] E. Freire, E. Ponce, F. Rodrigo, and F. Torres. Bifurcation sets of continuous piecewise linear systems with two zones. International Journal of Bifurcation and Chaos, 8:2073–2097, 1998.

[97] E. Freire, E. Ponce, and J. Ros. Limit cycle bifurcations from center in symmetric piecewise-linear systems. International Journal of Bifurcation and Chaos, 9:895–907, 1999.

[98] N. Arima, H. Okazaki, and H. Nakano. A generation mechanism of canards in a piecewise linear system. IEICE Transactions on Fundamentals of Electronics, Communications and Computer Sciences, E80:447–453, 1997.

[99] S. Coombes, R. Thul, and K. C. A. Wedgwood. Nonsmooth dynamics in spiking neuron models. Physica D, 241:2042–2057, 2012.

[100] M. Desroches, A. Guillamon, E. Ponce, R. Prohens, S. Rodrigues, and A. E. Teruel. Canards, folded nodes and mixed-mode oscillations in piecewise-linear slow-fast systems. SIAM Review, 58:653–691, 2016.

[101] M. Desroches, S. Fernández-García, and M. Krupa. Canards in a minimal piecewise-linear square-wave burster. Chaos, 26:073111, 2016.

[102] S. Fernández-García, M. Desroches, M. Krupa, and E. Teruel. Canard solutions in planar piecewise linear systems with three zones. Dynamical Systems: An International Journal, 31:173–197, 2015.

[103] S. Fernández-García, M. Desroches, M. Krupa, and F. Clément. A multiple timescale coupling of piecewise-linear oscillators. application to a neuroendocrine system. SIAM J. Appl. Dyn. Syst., 14:643–673, 2015.

[104] A. Roberts. Canard explosion and relaxation oscillation in planar, piecewise-smooth, continuous systems. SIAM J. Appl. Dyn. Syst., 609-624:15, 2016.

[105] A. Kaminaga, V. K. Vanag, and I. R. Epstein. A reaction-diffusion memory device. Angew. Chem. Int. Ed., 45:3087–3089, 2006.

[106] P. Goldman-Rakic. Cellular basis of working memory. Neuron, 14:477–485, 1995.

[107] K. Wimmer, D. Q. Nykamp, C. Constantinidis, and A. Compte. bump attractor dynamics in pre-frontal cortex explains behavioral precision in spatial working meory. Nature Neurosci., 17:431–441, 2014.

[108] M. Camperi and X.-J. Wang. A model of visuospatial working memory in prefrontal cortex: Recurrent network and cellular bistability. J. Comp. Neurosci., 5:383–405, 1995.

[109] K. Tornheim. Are metabolic oscillations responsible for normal oscillatory insulin secretion. Diabetes, 46:1375–1380, 1997.

[110] H. F. Chou, N. Berman, and E. Ipp. Oscillations of lactate released from islets of Langerhans: evidence for oscillatory glycolysis in beta-cells. Am J Physiol., 262:E800–805, 1992.

[111] R. Bertram, A. Sherman, and L. Satin. Metabolic and electrical oscillations: Partners in controlling pulsatile insulin secretion. Am. J. Physiol. Endocrinol. Metab., 293:890–900, 2007.

[112] M. J. Merrins, B. Fendler, M. Zhang, A. Sherman, R. Bertram, and L. Satin. Metabolic oscillations in pancreatic islets depend on the intracellular ca^2^+ level but not ca^2^+ oscillations. Biophys. J., 99:76–84, 2010.

[113] R. L. Burden and J. D. Faires. Numerical analysis. PWS Publishing Company - Boston, 1980.

[114] R. Kuske and T. Erneux. Localized synchronization of two coupled solid state lasers. Optics Communications, 139:125–131, 1997.

[115] R. Kuske and T. Erneux. Bifurcation to localized oscillations. Euro. J. Appl. Math., 8:389–402, 1997.

[116] H. G. Rotstein and R. Kuske. Localized and asynchronous patterns via canards in coupled calcium oscillators. Physica D, 215:46–61, 2006.

[117] Y. Kuramoto. Chemical Oscillations, Waves, and Turbulence. Sringer-Verlag, Berlin, 1984.

[118] D. Hansel, G. Mato, and C. Meunier. Clustering and slow switching in globally coupled phase oscillators. Phys. Rev. E, 48:3470–3477, 1993.

[119] P. Ashwin, G. Orozs, J. Wordsworth, and S. Townley. Dynamics on networks of cluster states for globally coupled phase oscillators. SIAM J. Appl. Dyn. Syst., 6:728–758, 2007.

[120] E. Brown, P. Holmes, and J. Moehlis. Globally coupled oscillator networks. In Perspectives and Problems in Nonlinear Science: A Celebratory Volume in Honor of Larry Sirovich, K. Sreenivasan, E. Kaplan, andJ. Marsden, eds., Springer, New York, pages 183–215, 2003.

[121] K. Y. Tsang, R. E. Mirollo, S. H. Strogatz, and K. Wiesenfeld. Dynamics of globally coupled oscillator arrays. Physica D, 48:102–112, 1991.

[122] S. H. Strogatz. From Kuramoto to Crawford: exploring the onset of synchronization in populations of coupled oscillators. Physica D, 143:1–20, 2000.

[123] A. Birzu and K. Krischer. Resonance tongues in a system of globally coupled oscillators with time-periodic coupling strength. Chaos, 20:043114, 2010.

[124] M. Somani, M. A. Liwuw, and D. Luss. Evolution and impact of temperature patterns during hydrogen oxidation on a Ni ring. Chem. Engg. Sci., 52:2331, 1997.

[125] C. G. Assisi, V. K. Jirsa, and J. A. Kelso. Synchrony and clustering in heterogeneous networks with global coupling and parameter dispersion. Phys. Rev. Lett., 94:018106, 2005.

[126] X. R. Sailer, V. Beato, L. Schimansky-Geier, and H. Engel. Noise-induced effects in excitable systems with local and global coupling. In Analysis and control of complex nonlinear processes in physics, chemistry and biology. L. Schimansky-Geier, B. Fiedler, J. Kurths, E. Scholl, eds. (World Scientific), pages 1–42, 2007.

